# Polygenic Adaptation has Impacted Multiple Anthropometric Traits

**DOI:** 10.1101/167551

**Authors:** Jeremy J. Berg, Xinjun Zhang, Graham Coop

## Abstract

Our understanding of the genetic basis of human adaptation is biased toward loci of large phenotypic effect. Genome wide association studies (GWAS) now enable the study of genetic adaptation in polygenic phenotypes. We test for polygenic adaptation among 187 world-wide human populations using polygenic scores constructed from GWAS of 34 complex traits. We identify signals of polygenic adaptation for anthropometric traits including height, infant head circumference (IHC), hip circumference and waist-to-hip ratio (WHR). Analysis of ancient DNA samples indicates that a north-south cline of height within Europe and and a west-east cline across Eurasia can be traced to selection for increased height in two late Pleistocene hunter gatherer populations living in western and west-central Eurasia. Our observation that IHC and WHR follow a latitudinal cline in Western Eurasia support the role of natural selection driving Bergmann’s Rule in humans, consistent with thermoregulatory adaptation in response to latitudinal temperature variation.

**Author’s Note on Failure to Replicate:** After this preprint was posted, the UK Biobank dataset was released, providing a new and open GWAS resource. When attempting to replicate the height selection results from this preprint using GWAS data from the UK Biobank, we discovered that we could not. In subsequent analyses, we determined that both the GIANT consortium height GWAS data, as well as another dataset that was used for replication, were impacted by stratification issues that created or at a minimum substantially inflated the height selection signals reported here. The results of this second investigation, written together with additional coauthors, have now been published (https://elifesciences.org/articles/39725 along with another paper by a separate group of authors, showing similar issues https://elifesciences.org/articles/39702). A preliminary investigation shows that the other non-height based results may suffer from similar issues. We stand by the theory and statistical methods reported in this paper, and the paper can be cited for these results. However, we have shown that the data on which the major empirical results were based are not sound, and so should be treated with caution until replicated.

## Main Text

Decades of research in anthropology have identified anthropometric traits that show evidence of biological adaptation to climatic conditions as humans spread around the world over the past hundred thousand years.^1, 2, 3^ However, it can be challenging to rule out non-heritable environmental factors,^4, 5^ as opposed to genetic variation, as the primary cause of these phenotypic differences.^6^ Even for phenotypes where there is some confidence that some of the phenotypic differences among populations are due in part to genetic differences, it is often hard to rule out genetic drift as an alterative explanation to selection.^7, 8, 9^ The development of population-genetic methods and genomic data resources during the last few decades has enabled the interrogation of adaptive hypotheses and has produced an expanding list of examples of plausible human adaptations.^10, 11^ However, such approaches are often inherently limited to detecting adaptation in genetically simple traits via large allele frequency changes at a small number of loci, whereas many adaptations likely involve highly polygenic traits and so are undetectable by most approaches.^12, 13^ Genome-wide association studies (GWAS) have now identified thousands of loci underlying the genetic basis of many complex traits.^14, 15, 16^ These studies oer an unprecedented opportunity to identify adaptation in recent human evolution by detecting subtle shifts in allele frequencies compounded over many GWAS loci.^17, 18, 19, 20, 21, 22, 23^

We conducted a broad screen for evidence of directional selection on variants that contribute to 34 polygenic traits by studying the distribution of their allele frequencies in a dataset of 187 human populations (2158 individuals across 161 populations from the Human Origins Panel^24^ and 2504 individuals across 26 populations of the 1000 Genomes phase 3 panel^25^), making use of prior large-scale GWAS for these traits (see Table S1). We divided the genome into 1700 non-overlapping and approximately independent linkage blocks^26^ and choose the SNP with the highest posterior probability of association within the block.^27, 28^ For each trait, we calculate a polygenic score for each population as a weighted sum of allele frequencies at each of these 1700 SNPs, with the GWAS effect sizes taken as the weights. Figure 1 shows the distribution of these scores for height across our population samples.

**Figure 1:**
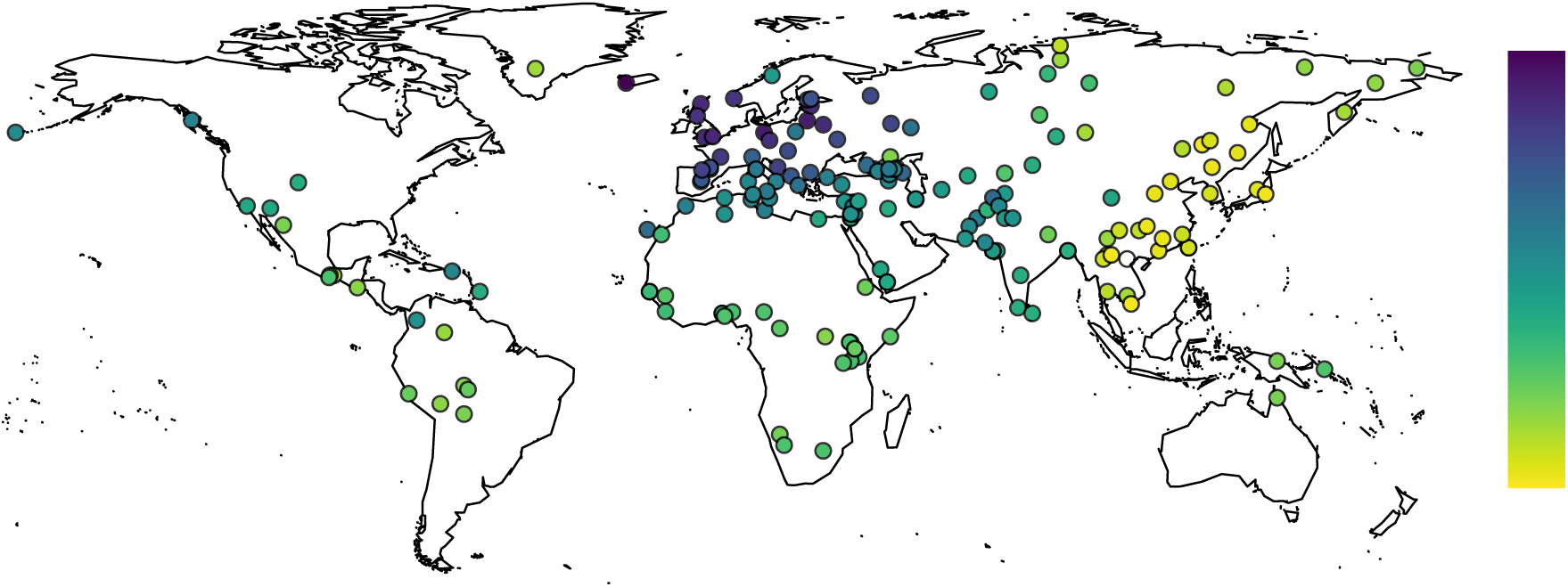
Polygenic Height Scores for 187 population samples (combined Human origin panel and 1000 genomes datasets), plotted on geographic coordinates. Blue corresponds to populations with the “tallest” polygenic height scores, and yellow the “shortest”.

These polygenic scores should not be viewed as phenotypic predictions across populations. For example, the Maasai and Biaka pygmy populations have similar polygenic scores despite having dramatic differences in height.^29^ Discrepancies between polygenic scores and actual phenotypes may be expected to occur either because of purely environmental influences on phenotype, or due to gene-by-gene and gene-by-environment interactions. We also expect that the accuracy of these scores when viewed as predictions should decay with genetic distance from Europe (where the GWAS were carried out), due to changes in the structure of linkage disequilibrium (LD) between causal variants and tag SNPs picked up in GWAS, and because GWAS are biased toward discovering intermediate frequency variants, which will explain more variance in the region they are mapped in than outside of it. These caveats notwithstanding, the distribution of polygenic scores across populations is informative about the history of natural selection on a given phenotype,^18^ and a number of striking patterns are visible in their distribution. For example, there is a strong gradient in polygenic height scores running from east to west across Eurasia (Figure 1)

To explore whether patterns observed in the polygenic scores were caused by natural selection, we tested whether the observed distribution of polygenic scores across populations could plausibly have been generated under a neutral model of genetic drift. To understand this null model, consider that a neutrally evolving allele has the same expected frequency across a set of independently evolving sub-populations. However, due to genetic drift, individual sub-populations will deviate from this expected frequency, with the variance of the sub-population frequencies given by F_ST_ *p* (1 − *p*), where p is the ancestral allele frequency, and F_ST_ is Wright’s “fixation index,”30 which can be measured from genome-wide data.^17, 31^ Our polygenic scores sum the contributions of a large number of effectively unlinked loci, which under our null model will experience genetic drift independently. It follows that under a model of genetic drift, the polygenic score of each of a set of independent sub-populations will be normally distributed, with variance of V AFST, where VA is the additive genetic variance of polygenic scores the ancestral population. Our test is based on a generalization of this simple relation in which we account for both variance and covariance among multiple populations that are non-independent due to common descent, migration, and admixture over the history of human evolution. Specifically, we model the joint distribution of polygenic scores as multivariate normal and use a generalized variance statistic (Q_X_) to measure the over-dispersion of polygenic scores relative to the neutral prediction, which is taken as evidence in favor of natural selection driving difference among populations in polygenic scores (see Methods and our previous study^18^ for details). Our approach is similar to classic tests of adaptation on phenotypes measured in common gardens, which rely on comparisons of the within and among-population additive genetic variance for phenotypes and neutral markers, i.e. Q_ST_/F_ST_ comparisons.32, 33, 34 Importantly, the neutral distribution we derive holds independent of whether the loci truly influence the trait in an additive manner (with respect to each other or the environment), and whether the GWAS loci are truly causal or merely imperfect tags. However, population structure in the original GWAS panels can confound signals of polygenic adaptation.^18, 20^ Modern methods are generally considered to be effective at controlling for the effects of population structure,^35^ and we proceed assuming that it has been adequately accounted for in the original GWAS panels.

We applied our test to each of the 34 traits across all populations, as well as within nine restricted regional groupings (Figure 2 and Table S3). Using our test across all populations as a general test for the impact of selection anywhere in the dataset, we find 5 signals of selection after controlling for multiple testing (p < 0.05/34). In each case of significant over-dispersion, the signal represents a small but systematic shift in allele frequency of a few percent across many loci, which would be undetectable by standard population-genetic tests for selection (see Table S6), such that the majority of the variance in polygenic scores is within populations as opposed to among populations (see Table S4). The traits involved include height, infant head circumference (IHC), hip circumference, waist-hip ratio (WHR), and type 2 diabetes (T2D). Although the sixth-strongest signal, waist circumference, failed to meet the multiple-testing correction, we include it in subsequent analyses due to its obvious relationship to WHR. We also found signals of selection on polygenic scores constructed for waist and hip circumference and waist-hip ratio when adjusted for BMI (Table S3), but we focus on the unadjusted versions for ease of interpretation. We do not replicate a previously reported signal of selection on BMI within Europe, but also note that the previous study used many more SNPs than we have in constructing polygenic scores, which likely explains the difference.^20^

The predominantly European ascertainment of GWAS loci can lead to apparent deviations from neutrality. Therefore all p values in Figure 2 and throughout the paper are derived from comparing test statistics against frequency-matched empirical controls, unless otherwise stated (see Text S1.3). This empirical matching is an important control. For example, the distribution of polygenic scores for Schizophrenia show a signal of over-dispersion under the naive null hypothesis, but not after controlling for the effects of ascertainment. More generally, the ascertainment and selection against disease phenotypes pose difficulties for the interpretation of tests of differentiation. Thus, although we see a signal of selection for decreased T2D polygenic scores in Europe, the interpretation of this signal likely requires the development of more explicit models of selection on disease traits (section S1.4).

### The Geography of Selection on Height

In addition to the known gradient of increased polygenic height scores in northern Europeans relative to southern Europeans (latitude correlation within Europe p = 6.3 × 10−^6^, see S2 and Methods for statistical details),^17, 18, 19, 20, 36^ we also find evidence that that natural selection has impacted polygenic height scores well outside of modern Europe. Polygenic scores decline sharply from west to east across Eurasia in a way that cannot be predicted by a neutral model (longitude correlation across Eurasia, p = 4.46 × 10−^15^; Figure 1), and they are overdispersed within each of our four population clusters (north, south/central, east, and west) across Asia, as well among Native Americans (Figure 2). Does this broadly Eurasian signal represents multiple independent episodes of selection on the genetic basis of height, or can it be explained by ancient selection on one or just a few populations, with modern signals reflecting variation in the extent to which modern populations derive ancestry from these ancient populations? For example, the signal of selection on height in East Asia is driven entirely by the Tu population sample, who have the highest polygenic height score among East Asian samples (p = 0.4329 for height in East Asia after the Tu are removed). Does this unusually high polygenic score reflect recent selection, or the fact that the Tu derive a proportion of their ancestry from an ∼800-year-old admixture event involving a population resembling modern Europeans^37^?

To test whether the height signal within Asia is due to a selective event shared with Europeans, we predicted the polygenic height scores across Asia given the deviation of European populations from the Asian mean, and each of the Asian sample’s genome-wide relationship to the European samples (see Figure 3, and Methods for details). We find that this prediction conditioned on Europeans are suffcient to explain most the divergence between the Tu and the other East Asian populations in our dataset (see sky blue dots in Figure 3), and eliminate the signal of selection among East Asian populations (p = 0.099 after conditioning). In fact, all signals of differential selection on height across Asia can be eliminated using these conditional predictions (p = 0.2019 after conditioning). This suggests that most of the selected divergence in our polygenic height scores across Eurasia can be attributed either to events which are predominantly ancestral to modern Europeans (but which have impacted other regions via admixture), or which lie along an early lineage which has contributed ancestry broadly across Eurasia.

**Figure 2:**
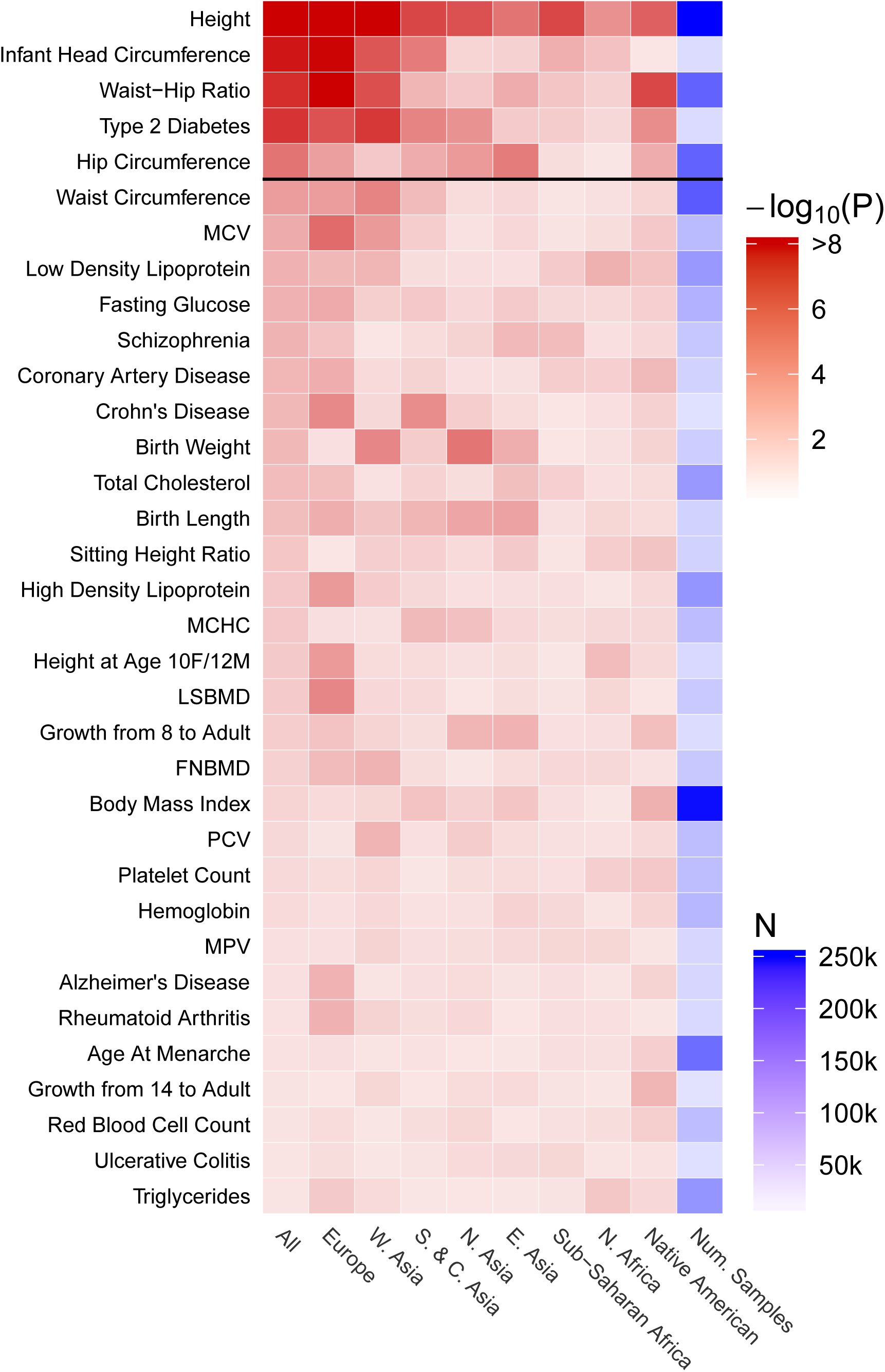
A heatmap showing the log10 p-values for the *Q*_*X*_ test statistic for over-dispersion of the polygenic scores for a trait among population samples. The ‘All’ column gives the p-value in the combined Human Origin and 1000 Genomes dataset. See S2 and S1 for the definitions of the regional groupings. Each subsequent column gives the score in each geographic subregion. MCV: Mean red blood cell volume; MCHC: Mean cell hemoglobin concentration; LSBMD: Lumbar spine bone mineral density; FNBMD: Femoral neck bone mineral density; PCV: Packed red blood cell volume; MPV: Mean platelet volume.

**Figure 3:**
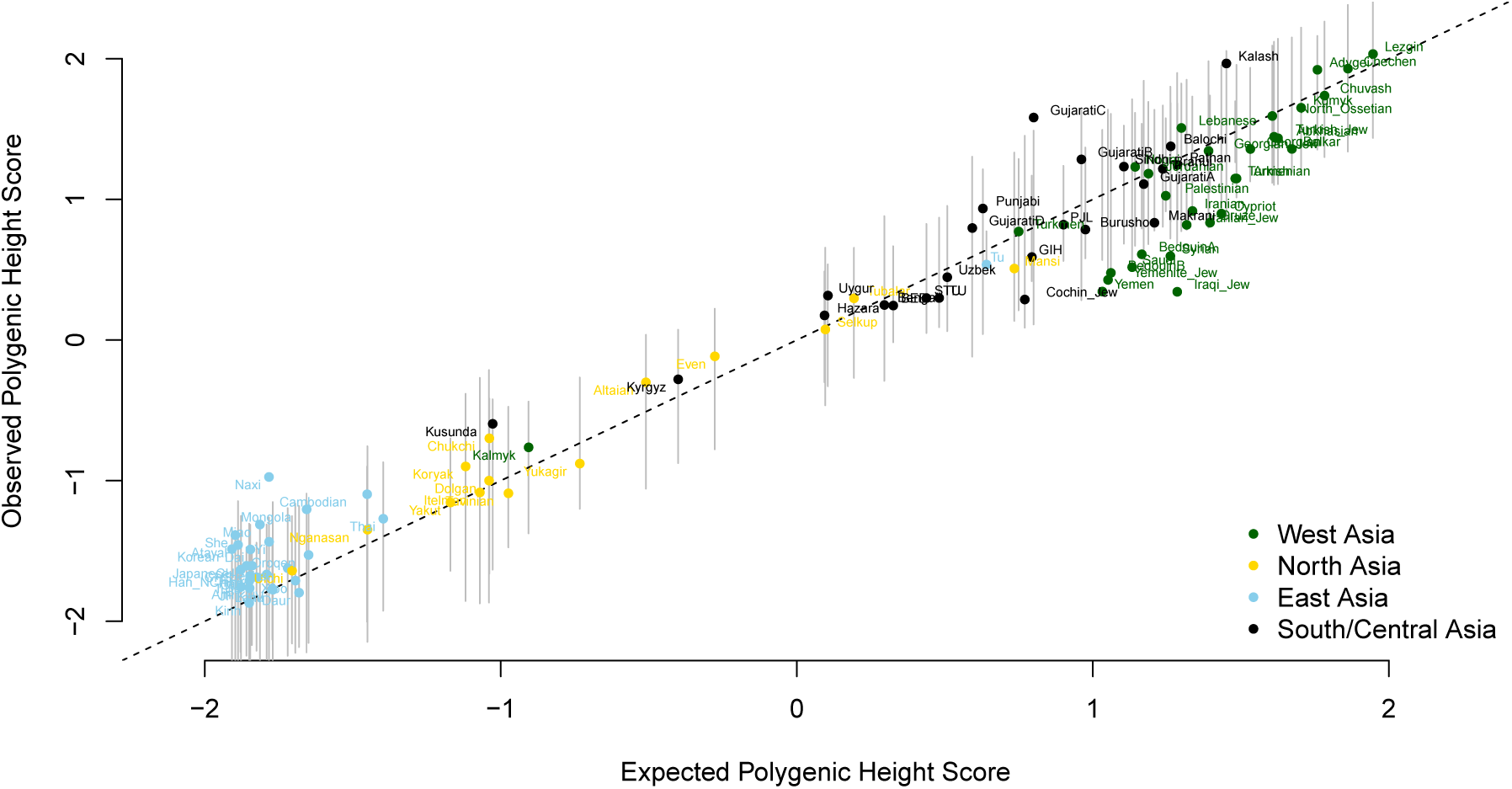
Polygenic height scores in Asia are well-predicted by a model conditioned on European height scores, consistent with selection occurring in a shared ancestral population. An individual population sample’s position along the x axis gives the genetic height score predicted on the basis of scores observed in Europe and their relatedness to the European samples, whereas their position along the y axis gives the true polygenic height score (see Methods for statistical details). The dashed line gives the one-to-one line along which all populations would fall if the predictions were perfectly accurate, whereas the vertical gray lines give population-specific 95% confidence intervals under genetic drift.

To gain further clarity about the history of selection on height, we examined polygenic height scores in a set of ancient DNA samples from Western Eurasia.^19, 38, 39^ In Figure 4A we plot estimates of the polygenic score through time for ancient and modern samples, and in Figure 4B a heatmap of signed p-values from our test of selection applied to pairs of populations (for more detail see Text S1.5). The earliest unambiguous signal of selection for increased height is found approximately 15,000 years ago in the Villabruna cluster of hunter-gatherers, who have significantly increased polygenic scores relative to earlier pleistocene hunter-gatherers (e.g. Villabruna vs Ust’-Ishim p = 0.0015, Villabruna vs Kostenki14 p = 0.0244, Villabruna vs Vestonice p = 0.003). The Mal’ta sample also appears to have an elevated polygenic score, on par with modern Europeans, but it is not significantly different from the earlier pleistocene hunter-gatherers in pairwise tests. Moving into the Holocene, the western, Scandanavian, and Caucasus hunter-gatherers (WHG, SHG, and CHG respectively) all have signficiantly increased polygenic height scores when compared to any of the early pleistocene hunter-gatherers. While WHG and SHG share a significant amount of ancestry with the Villabruna cluster, CHG do not, having separated approximately 46kya (along with Mal’ta and the Eastern hunter-gatherers: EHG) from the lineage leading to Villabruna/WHG.^40, 38^ Many ancient samples have ancestry nested within this split between Villabruna/WHG and CHG, but seemingly do not inherit a signal of selection for increased height (including pleistocene hunter-gatherers Kostenki14 and Vestonice^41, 38^). It is therefore unlikely that the signals we observe can be traced to a single selective event common to Villabruna/WHG/SHG and to CHG. Instead, our results are potentially consistent with at least two independent episodes of selection for increased height among pleistocene and/or holocene hunter-gatherers: at least one in the west, affecting Villabruna, WHG, and SHG, and one in the east, affecting CHG (and potentially Mal’ta).

**Figure 4:**
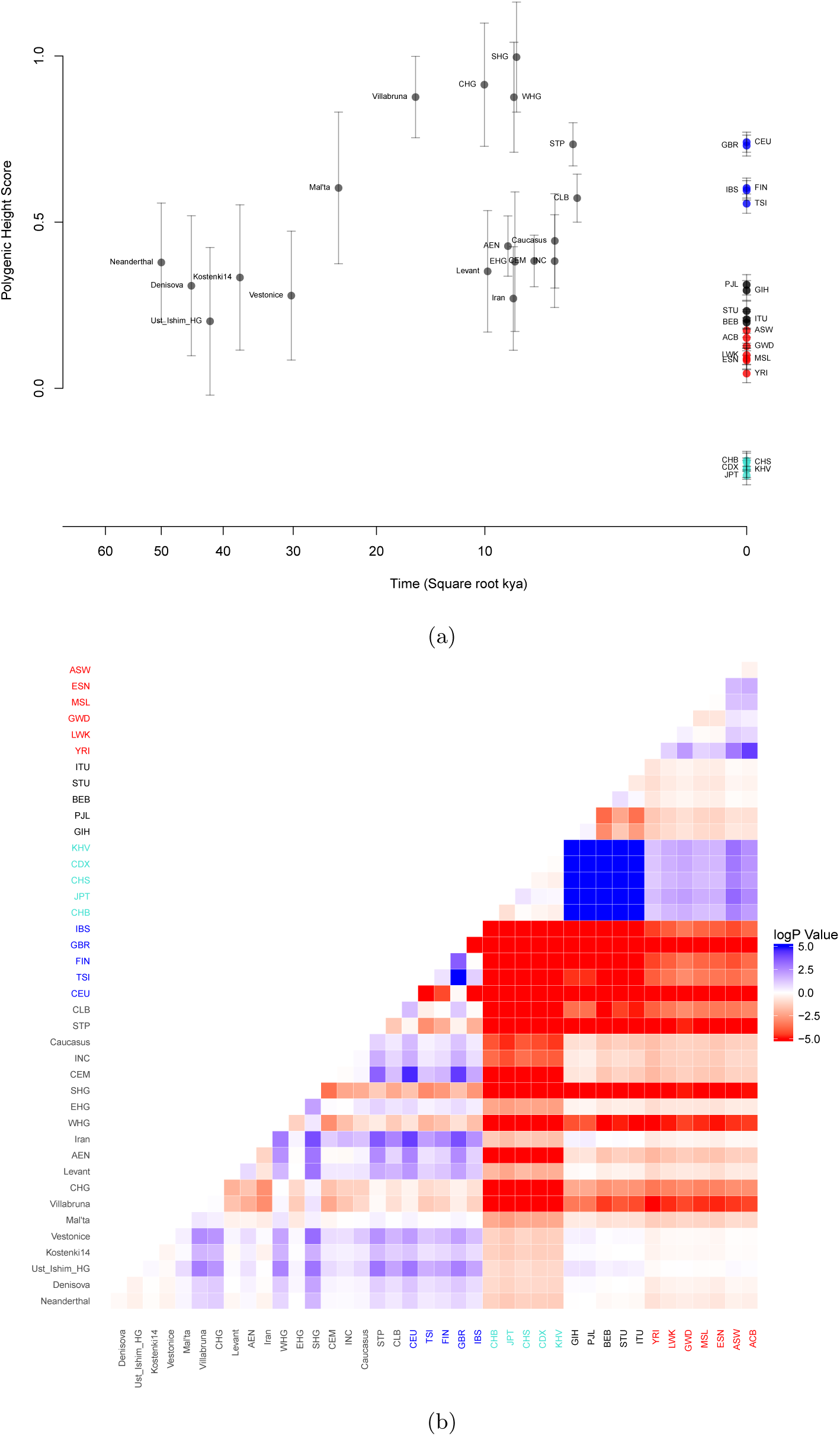
A) Polygenic height scores for ancient and the modern 1000 genomes population samples. Each dot show the mean polygenic score for the labeled sample, and the error bars give the 95% confidence interval. The x coordinate of each sample is positioned at the mean of the calBP dates for the samples, plotted using a square root transfor to help visualize the spread of ancient populations. AEN, Anatolian Neolithic; WHG, Western hunter-gatherer; CEM, central European Early and Middle Neolithic; INC, Iberian Neolithic and Chalcolithic; CLB, central European Late Neolithic and Bronze Age; STP, steppe. **B)** A heat map of log10 p-values for pairwise *Q*_*X*_ tests, the p-values are signed by the difference in polygenic score (shades of red denotes the row sample having higher polygenic score than the column sample, and blue the converse)

The Yamnaya-related steppe samples (STP) also show a signal of selection for increased polygenic height scores (e.g. STP–Ust’-Ishim p = 0.001, STP–Vestonice p = 0.004).^19, 42^ This signal is likely due to the fact that they draw ∼45% of their ancestry from a population related to the CHG,^19^ who they are not significantly different from (STP–CHG p = 0.62). In turn, the central European Late Neolithic and Bronze Age samples (CLB, including the Corded Ware and Bell Beaker culture) share the high polygenic height signal, and draw much of their ancestry from the expansion of the Yamnaya Steppe people.^43, 44^ In contrast, many of the European and Near East early Neolithic samples show little difference in scores relative to the early pleistocene hunter-gatherers and have significantly lower polygenic height scores than Villabruna/WHG/SHG and CHG samples and the populations with Yamnaya ancestry (e.g. Levant–SHG p = 0.001, Levant–CHG p = 0.01, Levant– STP p = 0.014). We do not find support for Mathieson and colleagues’^19^ suggestion of selection for reduced height in Iberian Neolithic samples relative to Anatolian Neolithic (p = 0.90, see also^42^).

Taken together, our results suggest that much of the variation we observe among modern Eurasian populations for polygenic height scores can be traced to variation in the amount of the WHG and Yamnaya/CHG ancestry they have inherited. For example, modern Europeans can be described approximately as a mixture between WHG, Yamnaya, and early Neolithic farmers from Anatolia,^43^ and the variation in the relative proportion of ancestry derived from these three sources explains a substantial amount of the variation in polygenic height scores (see Figure S10).^19, 42^ Similarly, Yamanaya/CHG ancestry decays from west to east across both northern and southern Asia,^40, 44^ consistent with the cline of decreasing polygenic height scores moving from west to east across the continent.

Finally, we note that we can reject neutrality in pairwise comparisons between modern East Asian populations and certain ancient samples that do not appear to be involved in the signal of selection for increased height in the west (e.g. CHB–EHG p = 0.004, CHB–Levant p = 0.014, Mal’ta–CHB p = 0.006). As these ancient populations are distantly related to one another, and show no other signals of selection on height, this may indicate that selection drove polygenic height scores down somewhere in the history of East Asians. However, the intepretation of this signal is complicated by the fact that we cannot completely exclude that polygenic height scores were selected up in these ancient populations. Clarifying this signal will likely require investigation via more explict models of human demographic history^23^ as well as the incorporation of height GWAS from East Asia.

### Selection on Body Shape Polygenic Scores

As four out of the next five strongest signals beyond height also represent anthropometric traits, we focus the remainder of our efforts on these phenotypes. Due to genetic correlations between traits, it is possible that signals of selection on two (or more) distinct phenotypes actually represent only a single episode of selection, where one trait responds indirectly to selection on the correlated trait. Because the genetic correlation with height varies among these phenotypes (hip circumference: r = 0.39, IHC: r = 0.268, waist circumference: r = 0.22, and WHR: r = −0.08),^45, 46^ we expect *a priori* that signals for more tightly correlated phenotypes are more likely due to a correlated response to selection on height, whereas for example the WHR signal is more likely to be independent.

To test whether the new signals we observe represent selective events distinguishable from the height signal, we developed a multi-trait extension to our null model based on the quantitative-genetic multivariate-selection model of Lande and Arnold^47^ (see Methods and Supplementary Text Section S1.6). We condition on the observed polygenic height scores, and test whether the signal of selection on a second trait is still significant after accounting for a genetic correlation with height (a non-significant p-value is consistent with a correlated response to selection on height). Applying this test to our entire panel of populations, we find that conditioning on height ablates much of the signal for hip circumference (p = 0.0186 compared to p = 1.12 × 10−^4^ when not conditioning on height), whereas signals in IHC (p = 1.11 × 10−^5^ vs p = 5.37 × 10−^8^) and WHR (p = 3.57 × 10−^8^ vs p = 3.38 × 10−^7^) are less affected. Restricting to European populations only, height is better able to explain hip circumference (p = 0.1152 vs p = 3.4 × 10−3), waist circumference (p = 0.0104 vs p = 2.63 × 10−^3^), and IHC (p = 5.1 × 10−^3^ vs p = 1.41 × 10−^8^) signals, while the signal of selection on WHR again remains strong even after conditioning on height (p = 1.92 × 10−^8^ vs p = 6.03 × 10−^10^). WHR is genetically correlated within populations with hip (r = 0.316) and waist circumference (r = 0.729), but not with IHC (r = 0.01).^45, 46^ Conditioning on WHR is suffcient to explain waist circumference (global p = 0.1523 vs p = 3 × 10−^3^, Europe p = 0.5178 vs p = 2.6 × 10−^3^), but signals in HIP, IHC, and height are all independent of WHR (see Table S4). Together, these results suggest that we can distinguish the action of natural selection along a minimum of two phenotypic dimensions (i.e. height and WHR, or unmeasured phenotypes closely correlated to them). The signal of selection observed for hip circumference is likely due at least in part to selection on height, and the waist circumference signal is probably due to selection on a combination of height and WHR (or closely correlated phenotypes; we provide additional evidence for this claim in supplement section S1.6.2). Whereas IHC shows some evidence of being influenced by selection on height, a correlated response to height seems not to fully explain this signal.

Signals of divergence for both IHC and WHR polygenic scores are confined mostly to Europe and West Asia. For both traits the null model gives a significantly improved fit to the data when conditioned on Europe to explain West Asia and similar when conditioning on West Asia to explain Europe (Table S5). This suggests that, as is the case for Eurasian height scores, a substantial fraction of the divergence in IHC and WHR polygenic scores among modern populations across western Eurasia reflects divergence among ancient populations and subsequent mixture rather than recent selection.

### Bergmann’s Rule and Thermoregulatory Adaptation

For both IHC and WHR, the selective signal in Western Eurasia can be captured in large part by strong, positive latitudinal clines (p = 3.16 × 10−^15^ for IHC and p = 3.16 × 10−^7^ for WHR; Figure 6). These clines in polygenic scores support independent phenotypic evidence for larger and wider bodies and rounder skulls at high latitudes,^48, 1, 49, 2, 50, 51, 3^ consistent with Bergmann’s Rule,^52, 53^ and add genetic support for a thermoregulatory hypothesis for morphological adaptation, whereby individuals in colder environments are thought to have adapted to improve heat conservation by decreasing their surface area to volume ratio.

A broad range of selective mechanisms have been proposed to act on height variation.^54^ Because we do not detect any signal of selection on age at menarche, we think it unlikely that the height signal represents a correlated response due to life-history mediated selection on age at reproductive maturity.^55^ It has also been suggested that selection on height may be explained as a thermoregulatory adaptation.^54^ However, because the surface area to volume ratio is approximately independent of height,^56, 2^ the effect of height SNPs on this ratio is mediated almost entirely through their effect on circumference (hip and/or waist; see section S1.8). Because the signal of selection on height cannot be explained by conditioning on hip and waist circumference, it seems that the thermoregulation hypothesis cannot fully explain the signal of selection on height.

A second eco-geographic rule relevant to height is Allen’s rule,^57^ which predicts relatively shorter limbs in colder environments, again consistent with adaptation on the basis of thermoregulation. In support of this, human populations in colder environments are observed to have proportionally shorter legs, compared to those in warmer environments.^49, 58^ However, we detect no signal of selection on polygenic scores for the ratio of sitting to standing height (SHR); a measure of leg length relative to total body height.^59^ Indeed, by combining our height SNPs with their effect on SHR, we find a strong signal that both increases in leg length and torso length underlie the selective signal on height from North to South within Europe, and from East to West across Eurasia (see S1.9). This again suggests that thermoregulatory concerns are unlikely to fully explain signals of selection on height.

## Discussion

The study of polygenic adaptation provides new avenues for the study of human evolution, and promises a new synthesis of physical anthropology and human genetics. Here, we undertake a broad scan for evidence of polygenic adaptive divergence among modern human populations, with body size and shape phenotypes providing most of our strongest signals. We show for the first time that it is possible to reject a neutral model of evolution at height associated loci in comparissons between populations outside of Europe. Using ancient DNA, we show that patterns seen across modern populations are consistent with two independent episodes of selection for increased height in pleistocene hunter-gatherer populations that lived in western and west-central Eurasia during or shortly after the last glacial maximum, and then distributed ancestry widely across the continent. We also provide evidence for adaptive divergence of IHC and WHR in western Eurasia, independent of selection on height, and show that signals of selection on hip and waist circumference can likely be explained as correlated responses to selection on height and WHR (or some other closely correlated phenotypes).

It is conspicuous that the signals of adaptive divergence that we detect are mostly localized to western Eurasia, even in cases where it seems implausible that observed phenotypic differences could have been generated under neutrality (e.g. Maasai vs Biaka pygmy). However, the fact that we do not detect departures from neutrality in such cases should not necessarily be taken as evidence against selection. We should expect to be better-powered to detect selective events in populations more closely related to Europeans for two reasons. First, changes in the structure of linkage disequilibrium (LD) across populations should lead GWAS variants to tag causal variation best in populations genetically close to the European-ancestry GWAS panels.^60^ Second, gene-by-environment and gene-by-gene interactions can lead to changes in the additive effects of individual loci among populations,^61^ and therefore in the way that they respond to selection on the phenotype. We expect that these difficulties can be overcome or mitigated in the future through a combination of well-powered GWAS in multiple populations of non-European ancestry, access to a wider array of ancient DNA samples, and improved frameworks for the interpretation of signals of polygenic adaptation.^23^

The existence of latitudinal trends in the polygenic scores for WHR and IHC support the notion that some of the clinal phenotypic variation in body shape typically thought to represent thermoregulatory adaptation can be attributed to genetic variation driven by selection, while the ability of simple models to unify signals across broad geographic regions again suggests that these patterns could have been generated by a limited number of selective events. Evidence for adaptation on the basis of specific environmental pressures is most convincing when multiple populations independently converge on the same phenotype in the face of the same environmental pressure, a pattern for which we currently lack evidence. Therefore, while our evidence is consistent with adaptation to temperature environments, alternative explanations (e.g. adaptation to diet) are plausible.

## 1 Methods

### 1.1 Population Genetics Datasets

We downloaded the 1000 genomes phase 3 release data from the 1000 genomes ftp portal.^25^ We also used data from the Human Origins fully public panel^24^ which was imputed from the 1000 Genomes phase 3 as reference, using the Michigan imputation server,^62^ and restricting to SNPs with an imputation quality score (in terms of predicted r^2^) of 0.8 or greater (pers. comm. Joe Pickrell). The original genotype data can be downloaded from the Reich lab website (https://reich.hms.harvard.edu/datasets).

This combined dataset represent samples from 2504 people from 26 populations in the 1000 Genomes dataset and 2158 people across 161 populations from the Human Origins dataset, for a total of 4662 samples from 187 populations (S2). For global analyses we include all 187 populations. In regional analyses we exclude populations with a significant recent (i.e. < 500 years) African/non-African admixture to avoid confounding admixture with signals of recent selection within regions (see S2 and S1 for the regions).

### 1.2 Selection of GWAS SNPs

We took public GWAS results for a set of traits^28^ and combined them with additional anthropometric traits from the GIANT consortium and a subset of Early Growth phenotypes contributed by EGG Consortium. Table S1 gives a full list of the traits included in this study and the relevant references. For each trait we selected a set of SNPs with which to construct our polygenic scores as follows. For each SNP, we calculated an approximate Bayes factor summarizing the evidence for association at that SNP via the method of Wakefield,^63^ following Pickrell *e*t al (2016)^28^ (see their supplementary note section 1.2.1). We then used a published set of 1700 non-overlapping linkage disequilibrium blocks^26^ to divide the genome, after which we selected the single SNP with the strongest approximate Bayes factor in favor of association within each block to carry forward for analyses.

### 1.3 Polygenic Scores and Null Model

Given a set of L SNPs associated with a trait (L ≈ 1700), we construct the vector of polygenic scores 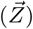 across all M = 187 populations by taking the sum of allele frequencies across the L sites (the vector 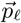 at site ℓ), weighting each allele’s frequency by its effect on the trait (αℓ) to give

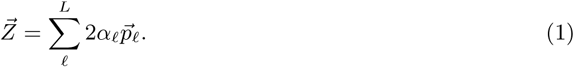

For each trait, we construct a null model for the joint distribution of polygenic scores across populations, assuming

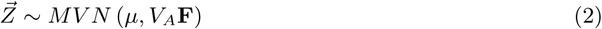

where 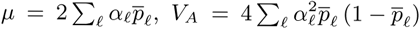. Here 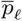 population samples (weighting all population samples equally), and **F** is the M × M population-level genetic covariance matrix.^18^ All polygenic scores are plotted in centered standardized form 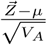. We use the Mahalanobis distance of 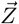 from its distribution under the null

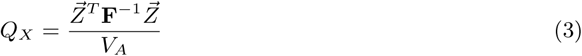

as a natural test statistic to assess the ability of the null model to explain the data (see Berg and Coop (2014)^18^ for an extended discussion). This test statistic should be X^2^ with M − 1 degrees of freedom under neutrality. However, in practice we are concerned that the ascertainment of GWAS loci may invalidate our null model, so we compare the test statistic to an empirical null (see Section S1.3)

### 1.4 Latitudinal and Longitudinal Correlations

We also test for selection-driven correlations between geographic variables (e.g. latitude) and a subset of our polygenic scores (see Berg and Coop (2014)^18^ and Section S1.1 for more details of the test). We take the standardized geographic variable and polygenic scores, and then rotate these vectors by the inverse Cholesky decomposition of the relatedness matrix **F**. These rotated vectors are in a reference frame where the populations represent independent contrasts under the neutral model. We take as our test statistic the covariance of these rotated vectors. We calculate the significance of the statistic by comparing to a null distribution generated by calculating null sets of polygenic scores assembled from resampled SNPs with derived frequency matched to the CEU population sample so as to mimic the effects of the GWAS ascertainment.

### 1.5 Analysis of Ancient DNA

We included a combined dataset of 63 Ancient Eurasian human population samples with date estimates from 45kya-2.5kya,^19, 38, 39^ combining these samples into pre-specified analysis clusters we took a set of 19 populations that had < 10% of height SNPs missing (see Table S7 for a list of ancient populations included). We compare these to the modern population samples from 1000 genome consortium data. We then took the subset of 724 of our 1700 height associated GWAS SNPs with low levels of missing data in these 19 ancient populations (6.2% averaged over populations).

Polygenic height scores were calculated as in Eq. (1), for loci with no counts in an ancient population we set to the frequency in the combined rest of sample. We construct the 95% credible intervals show in Figure 4A, by assuming that the the posterior of the underlying population frequency is independent across loci and populations and follows a beta distribution, with a uniform prior distribution, which is updated by our binomial sample of ancient counts. Using the variance of the posterior distribution at each locus, we then calculated the variance of the polygenic score (V_Z_), which follows from Eq. (1). The 95% credible-interval error bars in Figure 4A were then calculated as 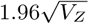 for each population.

For calculating Q_X_(eqn (3)) for pairs of population samples, we restricted the SNP set to the loci that had counts in both samples. Our p-values are calculated assuming that the pairwise Q_X_ statistic has a χ^2^ distribution, with one degree of freedom. We also constructed a null by flipping the signs of the GWAS effect of the loci at random, and found the χ^2^ p-values to be well callibrated.

### 1.6 Two-Trait Conditional Tests

Because some of the traits we examine are genetically correlated with one another, we were concerned that signals of selection observed for one trait might reflect a response to selection on another correlated trait. To determine whether genetic correlations might be responsible for some of our signals, we developed a multitrait extension to our neutral model that accounts for genetic covariance among traits. The extension is on the framework of Lande and Arnold.^47^

If 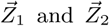 are vectors of polygenic scores for two different traits constructed according to equation (1), and the matrix 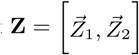 contains these vectors as columns, then under neutrality the distribution of **Z** is approximately matrix normal

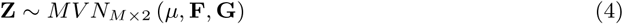

where the matrix µ contains the trait-specific means, **F** gives the population covariance structure among rows as in the single trait model, and **G** is the among trait additive genetic covariance matrix, the “G matrix” of multivariate quantitative genetics,^47^ estimated for a population ancestral to all populations in the sample. The diagonal elements of the 2 × 2 G matrix are given by the V_A_ parameters from above in the single trait model and the o-diagonal element (C_A,12_) corresponds to the additive genetic covariance between the two traits. Given this null model for the joint distribution of the two traits, we can construct a conditional model for the distribution of polygenic scores for trait 1, given the polygenic score observed for trait 2, as

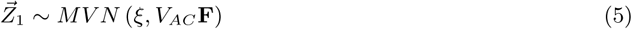

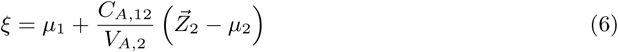

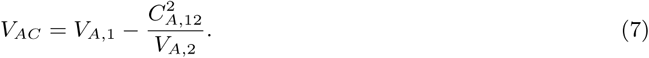

Given a value of C_A,12_ we can then use these conditional means and variances in equation (3) to form a conditional Q_X_ statistic and compare it to its null distribution. We take the failure to reject neutrality on the basis of the conditional Q_X_ statistic as consistent with the hypothesis that any response to selection observed for trait 1 is a result of selection on trait 2. Some of the traits we study have non-linear allometric relationships with each other, but because our polygenic scores are linear by construction our tests are robust to this non-linearity (see S1.7).

We experimented with estimating C_A,12_ on the basis of SNPs that overlap between the two traits in each genomic block. However, we were concerned about this approach to estimating genetic correlations not being a suffcient joint model for cases in which different SNPs within a block affected the two traits but were in linkage disequilibrium with one another, and therefore do not drift independently. To deal with this issue, we represent the genetic covariance among populations as

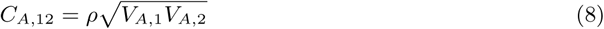

where ρ represents the genetic correlation between the two sets of polygenic scores. We pursued a conservative strategy, testing a range of values for ρ along a dense grid from −1 to 1 to ask whether *any* assumed genetic correlation between polygenic scores could plausibly allow one trait to be explained as a correlated response to another. As a further conservative measure, we allowed the genetic correlation used to calculate the conditional variance (Eq (7)) to be equal to zero, while allowing the ρ used to compute the conditional mean (Eq (6)) was not. This is a conservative approach, as it fits our conditional prediction to the mean, but allows the variance of the null model to remain as large as the unconditional model. The conditional two-trait p-values we present in the text, and the CI shown in two-trait Figure 5 and in the supplement, use this conservative approach. In practice our values of ρ are consistent with estimates of genetic correlations obtained from the LDscore approach,^45, 46^ given that our polygenic scores capture only a fraction of the total genetic variance for each trait.

**Figure 5:**
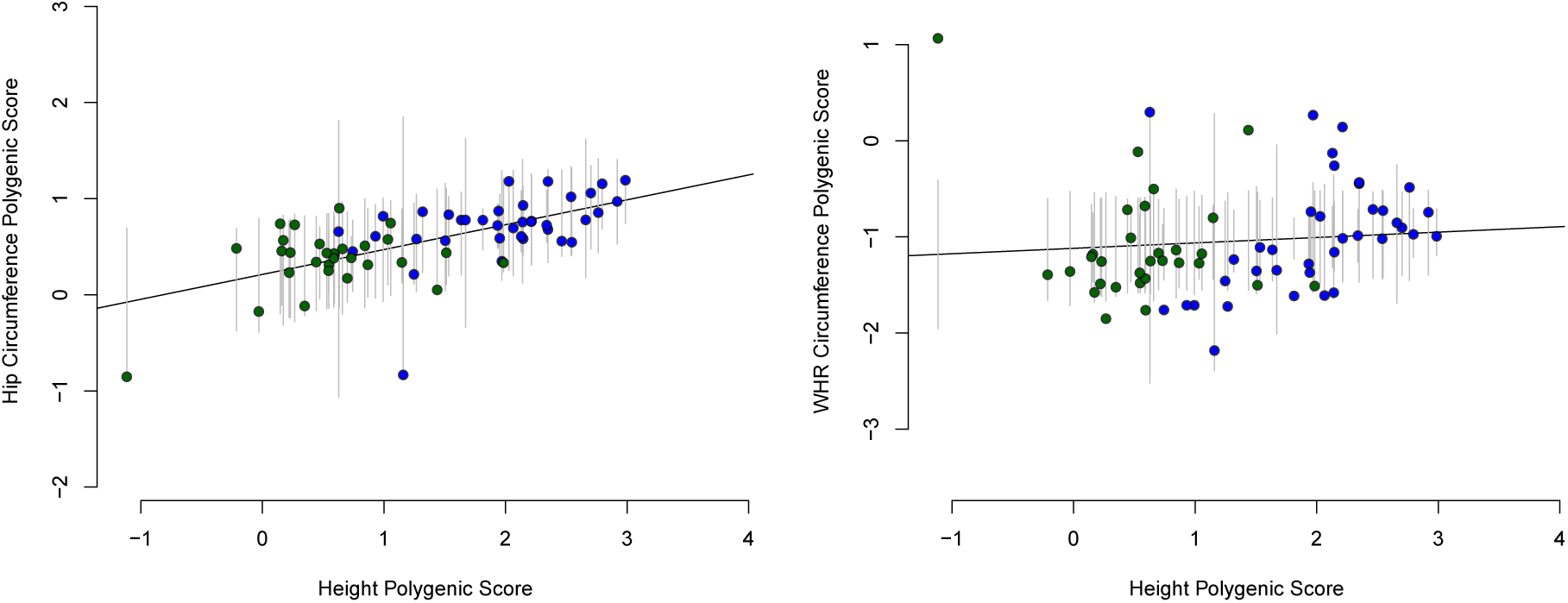
The overdispersion of genetic HIP scores among populations can be explained as a correlated response to selection on height, but such an effect cannot explain the signal of selection on the WHR polygenic scores. **A)** The observed polygenic HIP score (y axis) plotted against the height polygenic scores (x axis). We show only Western Eurasian population samples (blue dots: Europe; green dots: West Asia), as it is these samples which drive the majority of the signal. The line gives the best prediction for each sample’s polygenic HIP score according to the model of a correlated response to selection on height. Vertical lines give the 95% confidence interval of this prediction for each sample under this model. Most populations’ polygenic HIP scores lie within their confidence intervals, consistent with our failure to reject this conditional null model (main text). **B)** The same as A but now giving polygenic WHR scores rather than HIP. Note that for many populations the WHR scores lie outside of their 95% CI predictions based on genetic drift and correlated selection on height alone, consistent with the inability of this model to fully capture variation in polygenic WHR scores (main text)

**Figure 6:**
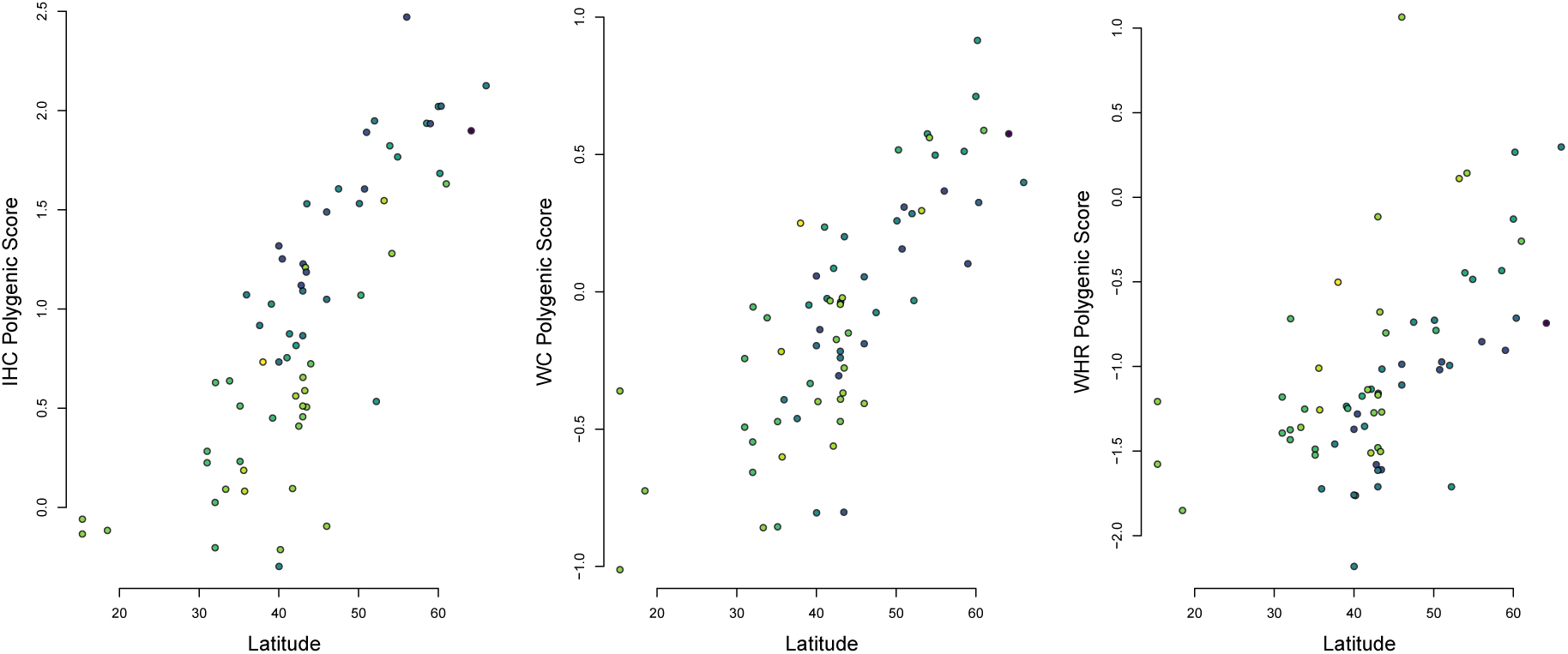
Genetic IHC, WC, and WHR score plotted against Latitude for the Western Eurasian population samples. The points are colored East to West (blue to yellow).

### 1.7 Single Trait Conditional Null Model

We also developed an extension of the null model for a single trait to test whether two (or more) signals of selection detected in different geographic regions might reflect a single ancestral event that occurred in an ancient population that has contributed ancestry broadly to modern populations.

Assume for example that we have detected a signal of selection among the population samples from region A (e.g. Europe) and among the population samples from (e.g. Asia), and we would like to test whether the signal detected in region B is due to a selective event that is also responsible for generating a signal of selection in region A. We first reorganize our samples into two blocks for the two regions

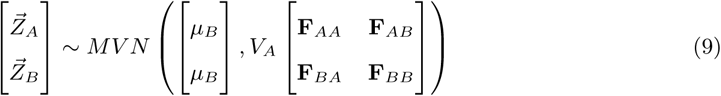

Where µ_B_ is the mean polygenic score in the set of populations being tested, the **F•,•**s refer to the sub-matrices of the relatedness matrix **F**, and **F** itself has been recentered at the mean of the test set (i.e. region B). Then the conditional distribution of polygenic scores in region B given the polygenic scores observed in region A is

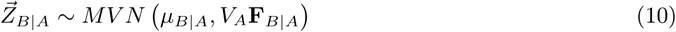

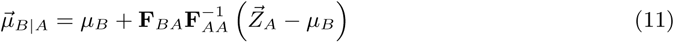

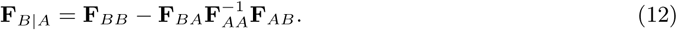

The conditional mean, 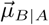 reflects the best predictions of population means in region B given the values observed in region A, whereas the conditional covariance matrix **F**_B|A_ reflects the scale and form of the variance around this expectation that arises from drift that is independent of drift in the ancestry of populations in region A.

We can then test for over-dispersion of polygenic score in region B 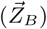 given the observed polygenic scores in region A by using 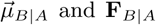 in (3) to construct a conditional Q_X_ score. We judge the statistical significance of this conditional Q_X_ score by comparing it to a frequency matched dataset, as with the standard test. We interpret a non-significant conditional Q_X_ score for region B as evidence that any selective signal of overdispersion in B is well explained by genome-wide allele-sharing with A. We view this as evidence that the selection signal in B overlaps that in A, due to selection in shared ancestral populations and admixture.

In Figure 3 we plot the observed polygenic scores for Asia against the predicted polygenic scores 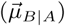 for Asia (B), conditional on the Europe population sample polygenic scores (A). The error bars are 95% CIs for each population sample, obtained from the variances on the diagonal of V_A_**F**_B|A_.

## Acknowledgements

We thank the Coop Lab and Doc Edge, Iain Mathieson, Emily Josephs, Joe Pickrell, Molly Przeworski, David Reich, Jeff Ross-Ibarra, Guy Sella, and Tim Weaver for helpful discussions and feedback on earlier drafts. The work was supported in part by an NSF GRFP (to JJB), the UC Davis Anthropology department (XZ), and NIGMS-NIH RO1 grants GM108779 to GC. JJB was also supported in part by R01 grants GM115889 to Guy Sella and GM121372 to Molly Przeworski.

## S1 Supplementary material

**Table S1:**
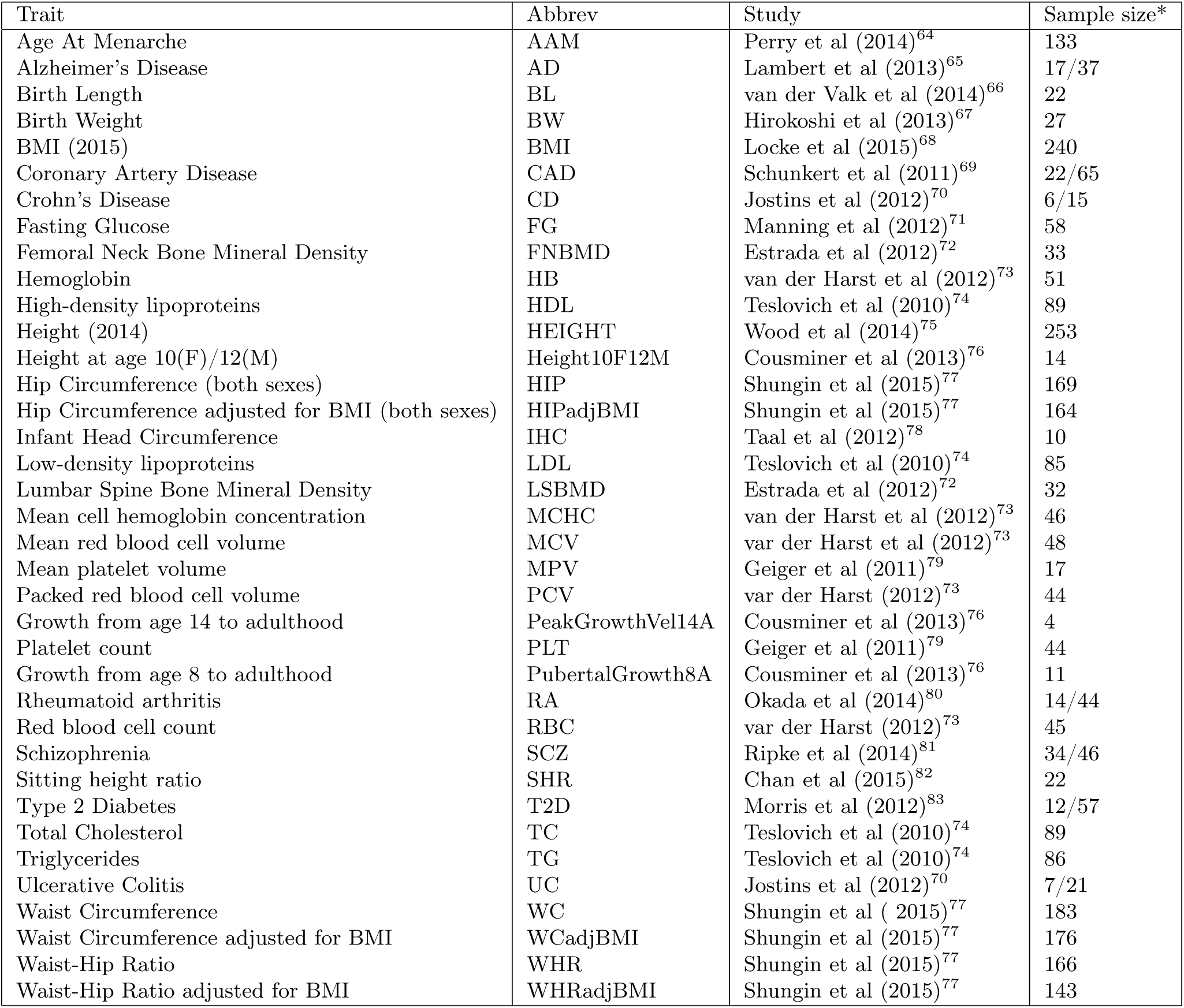
A list of all of the datasets tested (including those not directly mentioned in the main text), with citations for each study. *For case-control study sample sizes are given as Number of Cases/Number of Controls.

Table S2: A list of all population samples included in our analysis, along with the number of individuals per sample, and our geographic region assignment for each population.

Table S3: A table of the log10 p-values for the *Q_X_* test statistic for over-dispersion of the polygenic scores for a trait among population samples. The 'All' column gives the p-value in the combined Human Origin and 1000 Genomes dataset. See S2 and S1 for the regional definition for the definitions of the regional groupings. Each subsequent column gives the score in each geographic sub-region. MCV: Mean red blood cell volume; MCHC: Mean cell hemoglobin concentration; LSBMD: Lumbar spine bone mineral density; FNBMD: Femoral neck bone mineral density; PCV: Packed red blood cell volume; MPV: Mean platelet volume. Note that this table includes HIP, WC, and WHR adjusted for BMI, in addition to the 42 traits shown in Figure 2. These three additional traits were included to followup on the selection signals on HIP, WC, and WHR polygenic scores.

Table S4: Bivariate tests for evidence of correlated selection. Each row gives the results of a conditional *Q*_*X*_ test for evidence that a signal of selection in one trait can be explained as a correlated response to selection on another. Each row corresponds to a choice of two traits (one selected, one not), and a geographic region without which the test was run. The genetic correlation listed is that which gave the least significant p value (i.e. the most conservative test).

Table S5: Conditional region tests. Each row gives a particular combination of trait, test region, and conditioned region, and presents the *Q*_*X*_ statistics and associated p values for that test.

**Table S6:** Average allele frequency differences in the trait increasing allele between a few example populations. This table simply demonstrates that even for phenotypes with very strong differentiation at the polygenic value level, these differences are caused by relatively small average shifts spread across many loci

|  | CEU-CDX | TSI-CDX | YRI-CDX | TSI-CEU | YRI-CEU | YRI-TSI |
| --- | --- | --- | --- | --- | --- | --- |
| Height | 0.05728 | 0.04136 | 0.01136 | -0.01591 | -0.04591 | -0.03 |
| HIP | 0.02559 | 0.02137 | 0.02317 | -0.00422 | -0.00242 | 0.0018 |
| WC | 0.00689 | 0.00265 | 0.01357 | -0.00423 | 0.00668 | 0.01092 |
| WHR | -0.01993 | -0.02677 | -0.00357 | -0.00684 | 0.01635 | 0.0232 |
| IHC | 0.02831 | 0.01988 | -0.00635 | -0.00843 | -0.03466 | -0.02623 |
| T2D | -0.02939 | -0.02377 | 0.00138 | 0.00562 | 0.03077 | 0.02515 |

**Table S7:** A list of all of the ancient samples included in our analysis, along with the number of individuals per sample.

| Cluster | Full Name | Sample size (n) |
| --- | --- | --- |
| Afanasievo |  | 5 |
| Anatolia Neolithic |  | 24 |
| Andronovo |  | 3 |
| Armenia Chalcolithic |  | 5 |
| Bell Beaker LN | Bell Beaker Late Neolithic | 17 |
| Central LNBA | Central European Late Neolithic and Bronze Age | 35 |
| Central MN | Central European Middle Neolithic | 6 |
| CHG | Caucasus Hunter Gatherers | 2 |
| Hungary BA | Hungary Bronze Age | 12 |
| Hungary EN | Hungary Early Neolithic | 10 |
| Iberia EN | Iberia Early Neolithic | 4 |
| Iceman |  | 1 |
| LBK EN | Linearbandkeramik Early Neolithic | 15 |
| Poltavka |  | 7 |
| Sintashta |  | 5 |
| Srubnaya |  | 12 |
| WHG | Western European hunter-gatherers | 3 |
| Yamnaya Kalmykia |  | 6 |
| Yamnaya Samara |  | 9 |

Table S8: A table of all of the Pairwise *Q_X_* test statistics and p-values for the comparisons of the 19 ancient and modern Eurasian 1kg populations.

**Figure S1:**
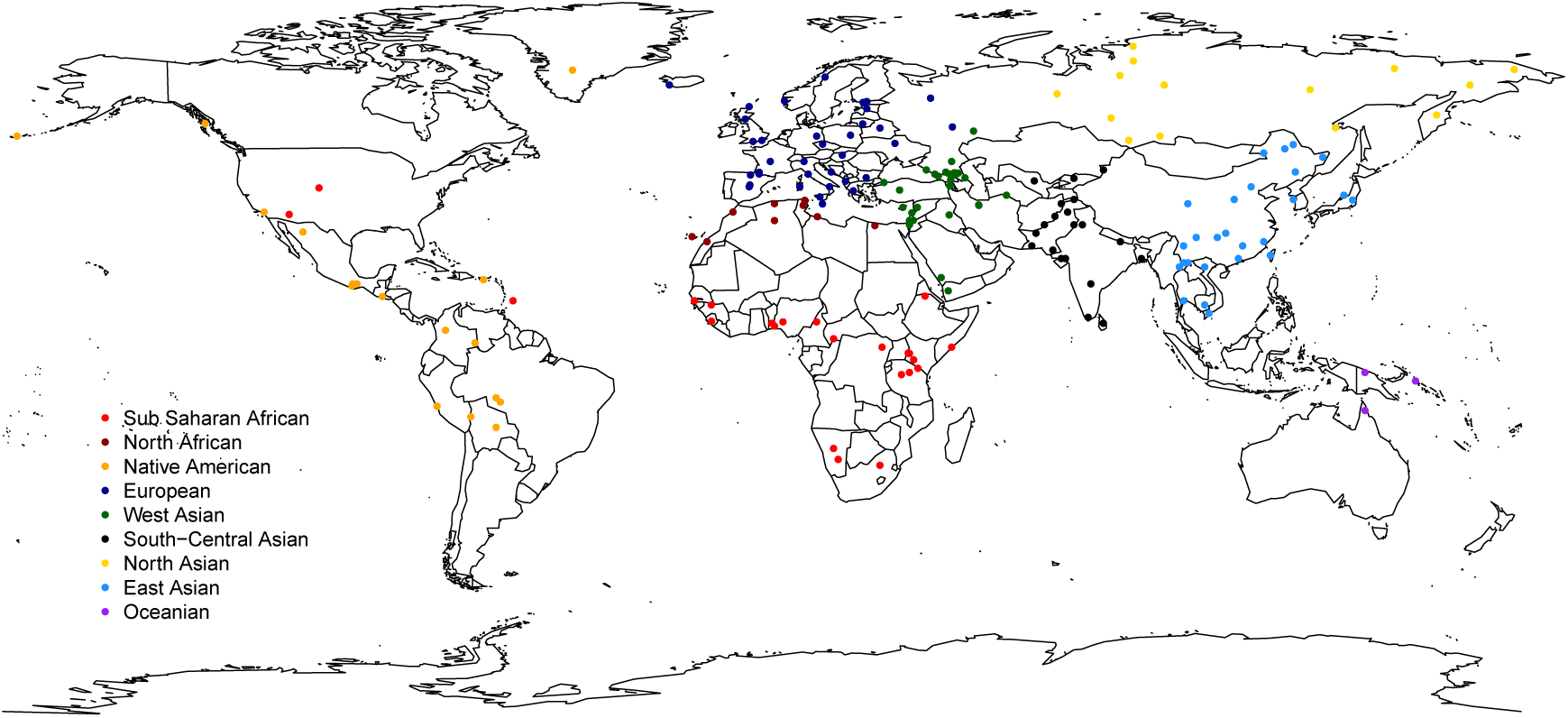
A map showing the locations of all 187 populations with each population colored according to a set of regional labels. Regional groupings were determined via a combination of geography and ancestry.

**Figure S2:**
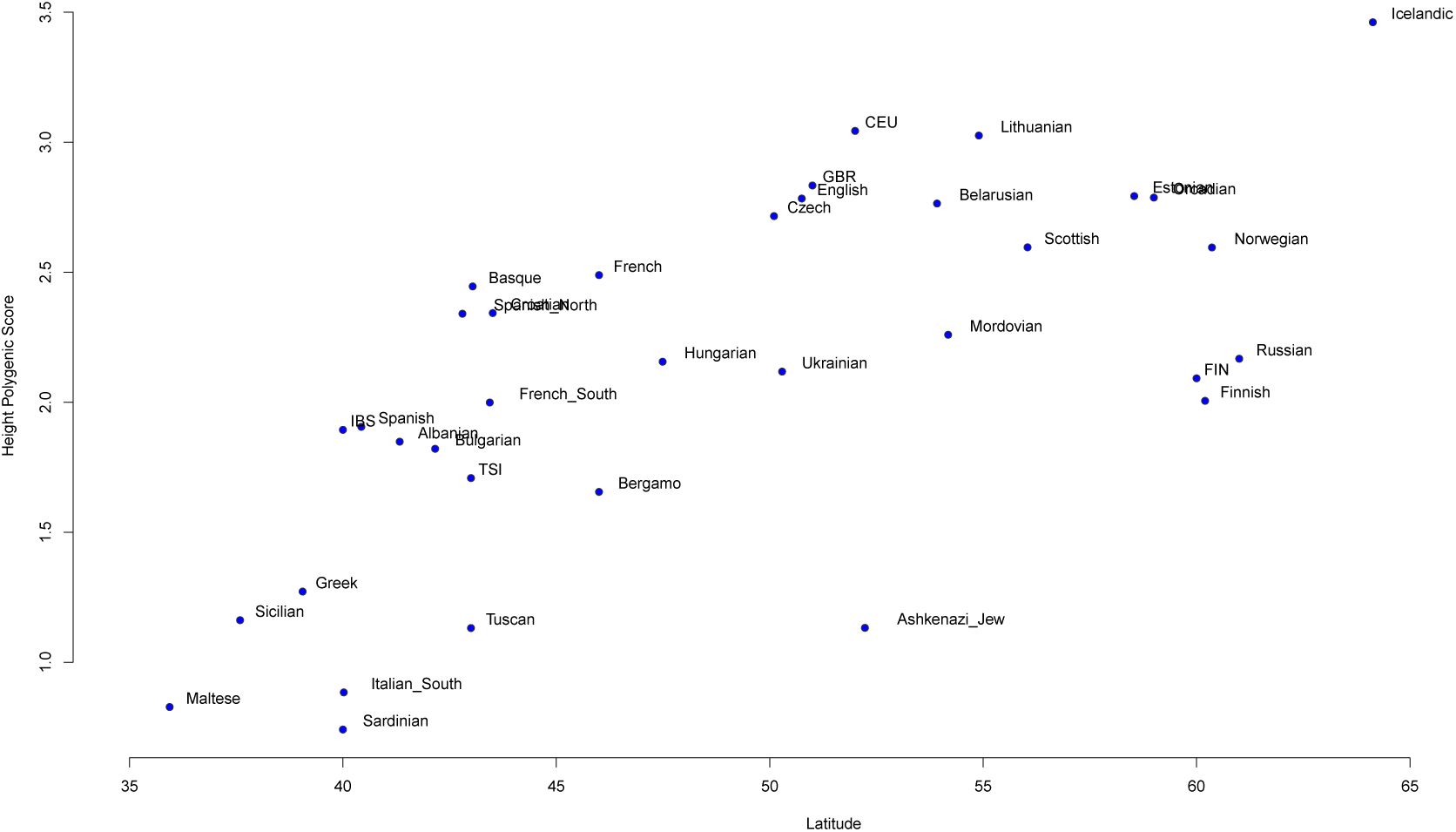
Polygenic scores for height within Europe plotted against latitude. This relationship is strongly significant even after controlling for population structure (*p* = 6.3 × 10^−6^), and represents our replication of previously reported latitudinal clines for height within Europe^17, 18, 19, 20, 36^

**Figure S3:**
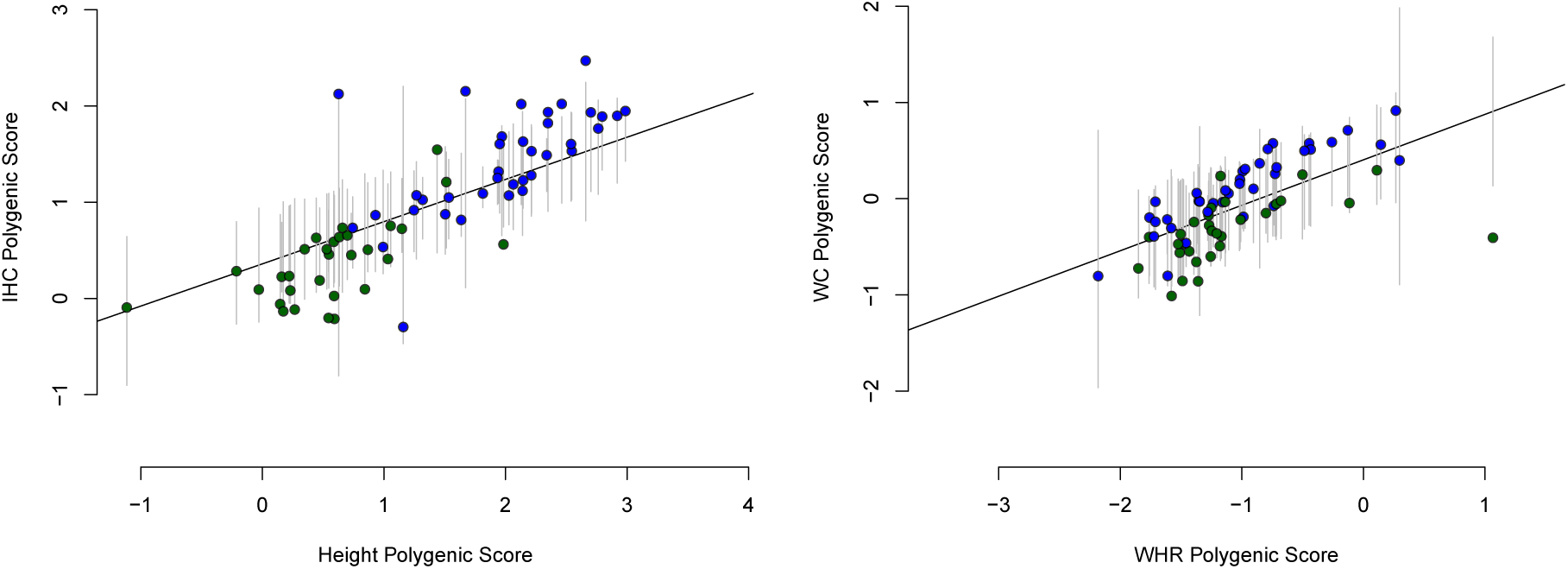
Left) Polygenic scores for IHC plotted against scores for height. Solid line gives the best prediction of IHC given height. Vertical grey lines give 95% confidence interval for each population. Note that a number of populations fall outside their error bars, consistent with the fact that we reject a neutral model for the evolution of IHC given height (see main text). Right) Same plot but using WHR to predict WC. Note that in this case, most populations fall well within their error bars, in line with the fact that WHR can adequately explain WC in our conditional *Q_X_* test.

**Figure S4:**
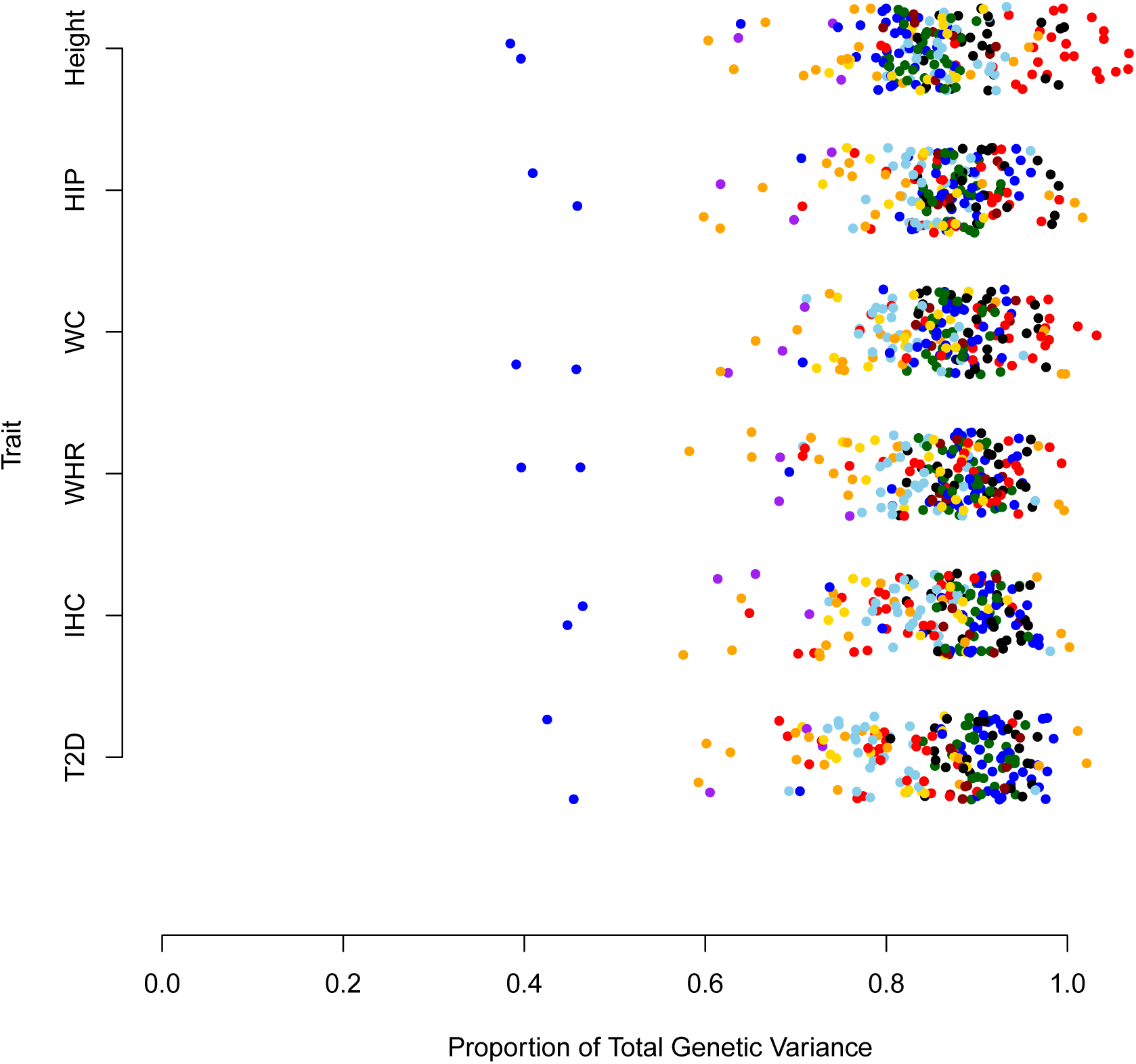
For each of the six traits showing evidence of selection, we show the proportion of total genetic variance for polygenic scores (based on Hardy-Weinberg and linkage equilibrium expectations) present within each population. Color scheme is the same as in Figure S1. Note that some of the variation among populations is due to differences in sample size, e.g. the two European (blue) populations with strongly reduced variance for each trait have only a single individual per sample.

### S1.1 Environmental Correlation Tests

We tested for unusually strong correlations between the polygenic scores for a given trait and an environmental or geographic (hereafter "environmental") variable as follows. Let 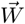 represent the vector of environmental variables recorded for each of the populations in our dataset. Now define 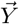 to be a mean centered and standardized version of 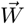

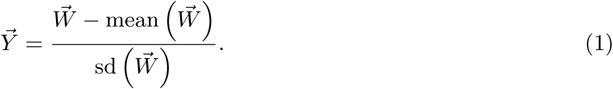

Next, recall that we have assumed that

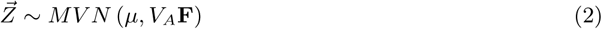

under the null model.

Now, let **C** be the Cholesky decomposition (or any other square root) of **F**, such that

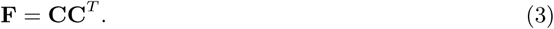

We next transform 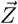

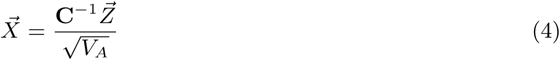

such that 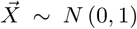 under the null (which each element of 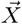 independent), but will retain information about any excess correlation with an environmental variable that is not predicted by drift and shared population history alone.

In order to test for such correlation, we must also transform the environmental variable

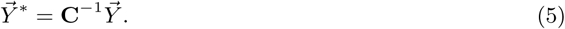

We then take the pearson product moment correlation between 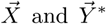 as a test statistic for the correlation between the polygenic scores and the environmental variable. Notably, because the test is performed in a rotated coordinate system that removes the effect of population structure, the test will have higher power and a lower false positive rate than a naive test of the untransformed polygenic scores. As with all of our other tests, in order to test for significance, we compare to a null distribution generated by calculating null sets of polygenic scores assembled from resampled SNPs matched for derived allele frequency to the CEU population sample so as to mimic the effects of the GWAS ascertainment.

### S1.2 Eigendecomposition of *Q*_*X*_ statistic

In constructing our empirical null statistic we make use of the fact that we can break the Q_X_ statistic down into the projection of the polygenic scores along each of the eigen-vectors of the matrix **F**.

Consider that our test statistic is given by

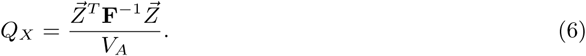

Now, we write the eigendecomposition of **F** as

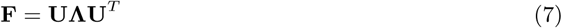

where **U** is a matrix containing the eigenvectors 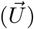 of **F** as its columns, and **Λ** is a matrix with the eigenvalues (λ) on the diagonal and zeroes elsewhere. For each eigenvector, we can define a statistic

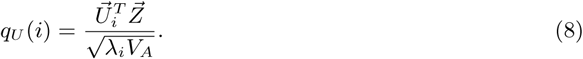

which is the slope of the regression of polygenic scores on the i^th^ eigenvector 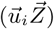 divided by the standard deviation of this regression coeffcient under the null 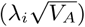 By the definitions of the multivariate normal distribution and the eigenvalue decomposition, this statistic has mean zero, a variance of one, and is linearly independent from all other such statistics q_u_(*j*) for *j* ≠ *i*. Note that the square of this statistic,

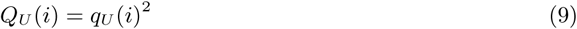

has a 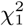 distribution under the null hypothesis, and because of their independence, our global test statistic Q_X_ can be written as a sum of the Q_u_(*i*)s:

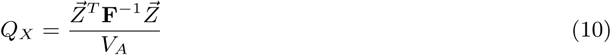

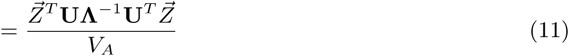

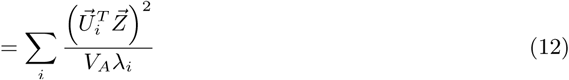

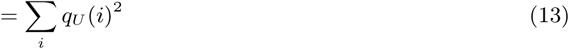

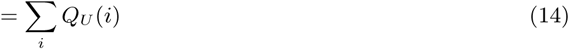

### S1.3 Empirical Null

The MVN model of drift is generally justified by supposing that we are told the frequency of an allele in a given population, and then asked to predict the joint distribution of allele frequencies across multiple descendant populations after a relatively small amount of neutral evolution. In this case, the MVN model is generally a good approximation to the diffusion (see previous work^31, 84, 18^ for more extended descriptions of this approximation).

However, consider an alternative case where instead of being told the frequency of the allele in the ancestral population, we are told the frequency in a single one of the descendant populations, and we are also told that the allele has a single mutational origin and we are told which allele is derived and which is ancestral. This knowledge alone is suffcient to violate the assumptions of the MVN model, as it must be the case that, looking backward in time from the present, the frequency of the derived allele decreases on average back until we reach the mutation which created it. Playing the tape forward in time, it is then clear that the expected change in allele frequency along the lineage leading to the conditioned upon population is not zero. Indeed, the effect is essentially a form of the "fictitious selection" described by Zhao and colleagues,^85^ that arises for alleles (neutral or otherwise) whose fate (forward or backward in time) is conditioned on.

Our case more closely resembles the latter example, as GWAS loci are ascertained in a particular present day population, and they must be at suffciently intermediate frequencies in order to be detected. We might therefore fear that the MVN does not strictly hold. Because positive signals in our test are generally created when the sign of an allele's effect on the trait is predictive of its distribution among populations (see previous work by Kremer and Le Corre^86, 87^ and ourselves^18^ for more extended discussions of this fact), we are most concerned about this problem when there is a correlation (within our set of GWAS loci) between the sign of an allele's effect on the trait and whether or not it is derived or ancestral.

In Figure S5, we show the the q_U_ (i) statistic for the schizophrenia dataset for the top 30 eigen-vectors of the population genetic covariance matrix (black dots), compared with an empirical null distribution for each q_U_ (i) statistic that is constructed by resampling SNPs which matched the derived allele frequency of the true associations in the 1000 genomes CEU panel (which we take as a proxy for the GWAS population). The exact procedure is detailed at the end of this section. Strikingly, we observe that for some of the eigenvectors (e.g. 1,2 and 4), the ascertainment matched empirical null distribution has a mean that is shifted away from zero, and toward the direction of the observed true q_U_ (i) statistic for that eigenvector.

This demonstrates that at least some of the signal we naively observe for schizophrenia has been created by the GWAS ascertainment procedure, and does not actually reflect the action of natural selection. Fortunately, these eigenvector statistics (i.e. the q_U_ (i)) offer an attractive route to controlling for these ascertainment effects. We define a recentered and standardized version of these eigenvector statistics as

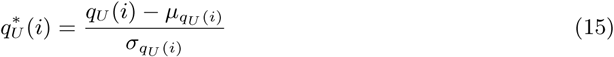

where µ_q*U* (i)_ is the mean of the empirically resampled null q_U_ (*i*) statistics, and σ_q*U (i)*_ their standard deviation. This ensures that each *q*_*U*_ (*i*) has mean zero and standard deviation one under the empirically recalibrated null, and we then take as our global test statistic the rescaled Q_X_ statistic

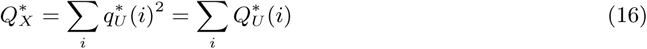

which should follow the appropriate χ^2^ distribution under the empirically calibrated null hypothesis. Throughout the paper we report p values derived from this empirical null unless otherwise stated.

**Figure S5:**
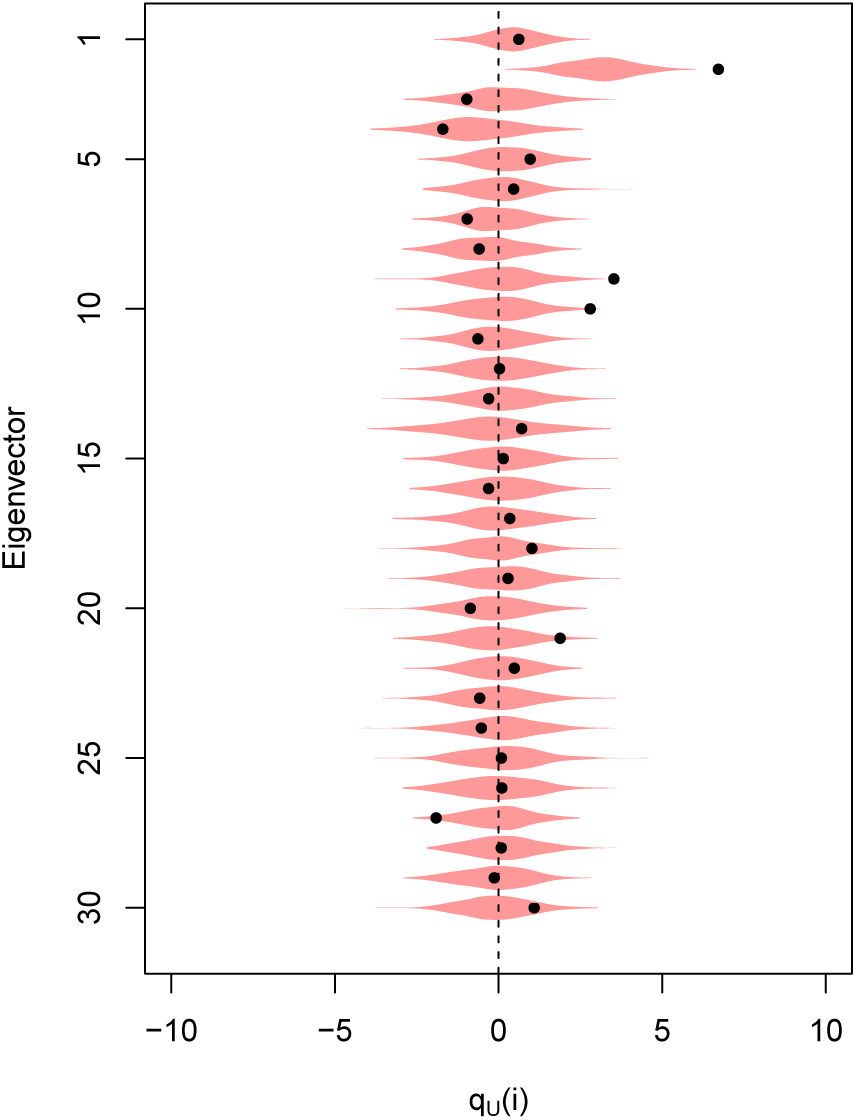
Violin plot of the eigenvector statistics for Schizophrenia. Each violin shows the empirical frequency matched null distribution for each eigenvector. Black dots give the eigenvector statistics for the true data. Violins which are not centered at zero, or which have variance greater than 1 indicate departures from the neutral null model caused by ascertainment

The two phenotypes most strongly impacted by this ascertainment effect are schizophrenia and type 2 diabetes. For schizophrenia, we find that after applying the recalibrated null the naive p value (p = 9.5 × 10^−6^) is reduced substantially (p = 0.018), such that in the end the strength of evidence against the null is relatively marginal. For type 2 diabetes, on the other hand, while comparison to the empirical null does reduce the strength of the signal, we still see significant evidence against the null even after comparison to the recalibrated null (naive p = 6.17 × 10^−11^ vs recalibrated p = 1.27 × 10^−6^).

### S1.4 Comparison between risk and protective variants for schizophrenia and type 2 diabetes

We noticed that the two phenotypes for which the empirically calibrated null deviated most strongly from the naive null were both disease traits (i.e. schizophrenia and type 2 diabetes). This is noteworthy because the loci identified for these phenotypes have been ascertained under the case-control study design, which is known to result in asymmetries in statistical power when the number of cases and controls are not equal (as they seldom are), with lower power to identify low frequency protective alleles than low frequency risk alleles.^88^ We were also concerned that a neutral model may not be an appropriate null for these phenotypes, as being diseases with severe fitness consequences, we might expect *a priori* that there would be systematic selection against risk increasing alleles and for risk decreasing alleles. Further, the combination of ascertainment bias and systematic selection against disease alleles might generate signals under our test that are real, in the sense that they represent a real long term response to selection on the GWAS loci identified, but may be misleading if interpreted naively as a signal of divergent selection among populations.

To understand how these two effects may have played a role in generating the signals we observe for SCZ and T2D, we separated alleles into two classes on the basis of whether the derived allele is a risk variant or a protective variant. We can then develop qualitative expectations about what sort of patterns we expect to see if either of the above mechanisms (asymmetric power and/or systematic selection against disease) are at play. Asymmetric power should have two major consequences. First, increased relative power to detect low frequency risk variants means that on average the most significant variant in a given block should be a derived risk variant greater than 50% of the time (as on average the rarer of the two alleles at a site will be the derived allele the majority of the time). This is consistent with what we observe for both diseases (T2D: 772 out of 1670 derived variants are protective, two tailed binomial test p = 0.002213; SCZ: 699 out of 1496 derived variants are protective, binomial test p = 0.012121). Second, protective variants that are detected should be systematically closer to 50% frequency than risk variants. Using the CEU as a proxy for the GWAS sample of primarily north-western European ancestry, we observe this for both diseases (T2D: derived protective mean frequency = 35.5% while derived risk mean frequency = 27.5%; SCZ: derived protective mean frequency = 35.5%, while derived risk mean frequency = 22.6%). This asymmetry alone would be suffcient to generate signal in our naive test prior to empirical calibration of the null model, and in our framework would be expected to present as selection for decreased risk in Europeans relative to other populations because derived allele frequencies should be lower in populations genetically distant to the European GWAS population. One way to think about this is that the "fictitious selection" described above due to conditioning in the GWAS is stronger for derived protective variants than for derived risk variants. Other populations' polygenic scores should also show this effect in a manner than depends upon how recently they share ancestry with the population in which the GWAS was done. However, because this effect involves no *actual* selection, it can be entirely controlled for by recalibration of the empirical null model to condition on the set of derived allele frequencies (as described in our Empirical Null Section).

In Figures S6 and S7 we show the observed mean frequency of derived risk and protective alleles in the CEU and YRI population samples. We also show the mean frequency for control derived alleles with matched frequencies in CEU. The lower frequency of the matched derived alleles in YRI clearly shows why a frequency matched null is necessary, as both the matched control protective alleles and the risk alleles have a higher average allele frequency in the CEU than the YRI due to the fictitious selection effect described above. However, both disease traits show some deviation away from the null expectation in these figures, particularly T2D, consistent with the fact that we strongly reject the neutral null model for T2D, but find only marginal signal at best for SCZ after controlling for ascertainment effects via our empirical null.

Our empirical null is based on neutral evolution, whereas we might expect these disease phenotypes to have been selected against *a priori* and in a similar manner across all populations, suggesting that our neutral model may not be an appropriate null. Adjusting our null expectation to account for the fact that we expect diseases to be systematically selected against is more challenging, and a detailed quantitative understanding is beyond the scope of this paper, but here we develop some qualitative expectations. First, consider a disease trait under constant negative selection, and compare protective alleles ascertained in the CEU (or a closely related sample of European individuals), to loci randomly sampled so as to have the same frequency distribution within the CEU. We expect that both the protective alleles and the control alleles should have a higher average allele frequency in the CEU than the YRI due to the fictitious selection effect described above. However, we might expect relatively little difference between protective and control alleles in YRI, as the fact that the protective alleles experience positive selection in the lineage leading to YRI while the controls do not is offset by the fact, given that they have been under positive selection, protective alleles were likely at lower frequency at the time of the split between the ancestors of CEU and YRI. Risk alleles, on the other hand, will have experienced the same fictitious selection in the lineage leading to CEU, but will have been held at lower frequency in the YRI due to selection against the disease. These two patterns are pictured together in Figure S8A.

However, our results for T2D (and to a lesser extent SCZ) do not match this qualitative expectation from a model of constant selection against the disease. It may be consistent with a model where selection pressures against the disease differ among populations. In the case of differential selection for lower population risk in CEU relative to YRI, we expect a different pattern, where protective alleles should be at systematically lower frequencies in YRI than their matched controls (reflecting their selection upward in frequency within CEU since the two populations split), while risk increasing alleles will tend to be at higher frequencies in YRI than matched controls, reflecting stronger negative selection against these alleles in the ancestors of the CEU than those of the YRI. This pattern is depicted in Figure S8B, and closely resembles that observed for T2D. However, a model based framework accounting for both ascertainment and purifying selection against disease will be needed to more fully demonstrate this.

### S1.5 Polygenic Height Scores and Ancient DNA

Overall, the previously detected selection signal of increased polygenic height scores in modern Northern Europeans is replicated in the ancestral Western Eurasian populations, with modern East Asians having the lowest polygenic height scores. This suggests that this signal of genetic height differentiation across Eurasia is old. The Anatolian Neolithic group and Iberian early farmers (Iberia EN, Iberia MN etc.) both show significantly reduced polygenic height scores relative to modern Europeans, that are only somewhat higher than East Asians. The Steppe populations (Andronovo, Srubnaya etc.) shows increased polygenic height scores, which are consistent with signals reported from Mathieson *e*t al.,^19^ with one exception for Hunter-Gatherer groups, which in our analysis show a distinct (and strongly statistically significant) increase in polygenic height scores. This increased signal^19^ in the Hunter-Gatherer groups likely reflects the increase in number of height associated loci included (724 vs. 180). In total, this is consistent with the view that the polygenic height score difference we see across Eurasia is old, and that as hypothesized by Mathieson *et al.*^19^ the modern gradient in polygenic height scores *within* Europe may be mostly driven by the mixture between

**Figure S6:**
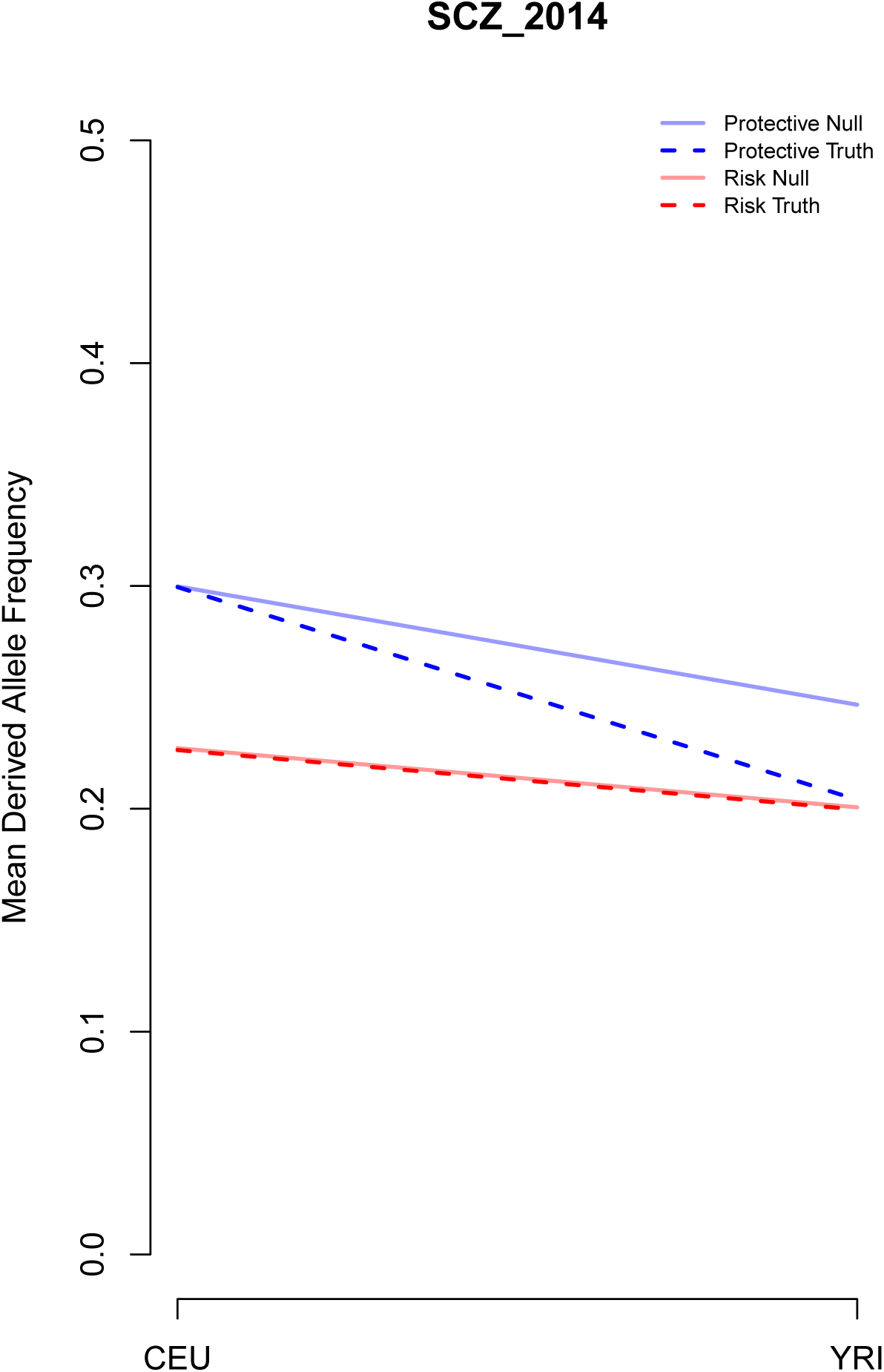
Observed pattern of SCZ risk and protective variants when comparing allele frequencies in CEU and YRI population samples.

**Figure S7:**
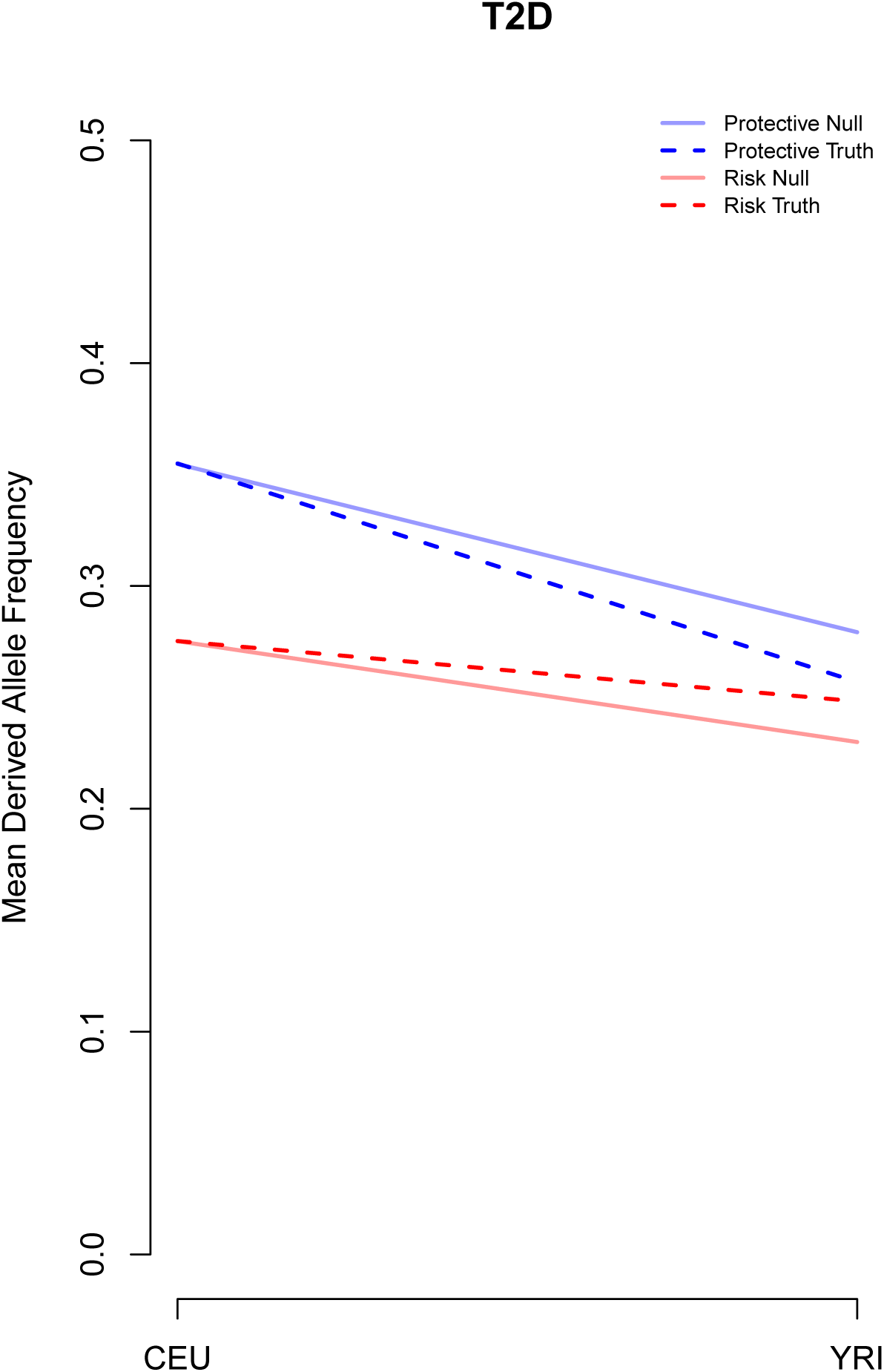
Observed pattern of T2D risk and protective variants when comparing allele frequencies in CEU and YRI population samples.

**Figure S8:**
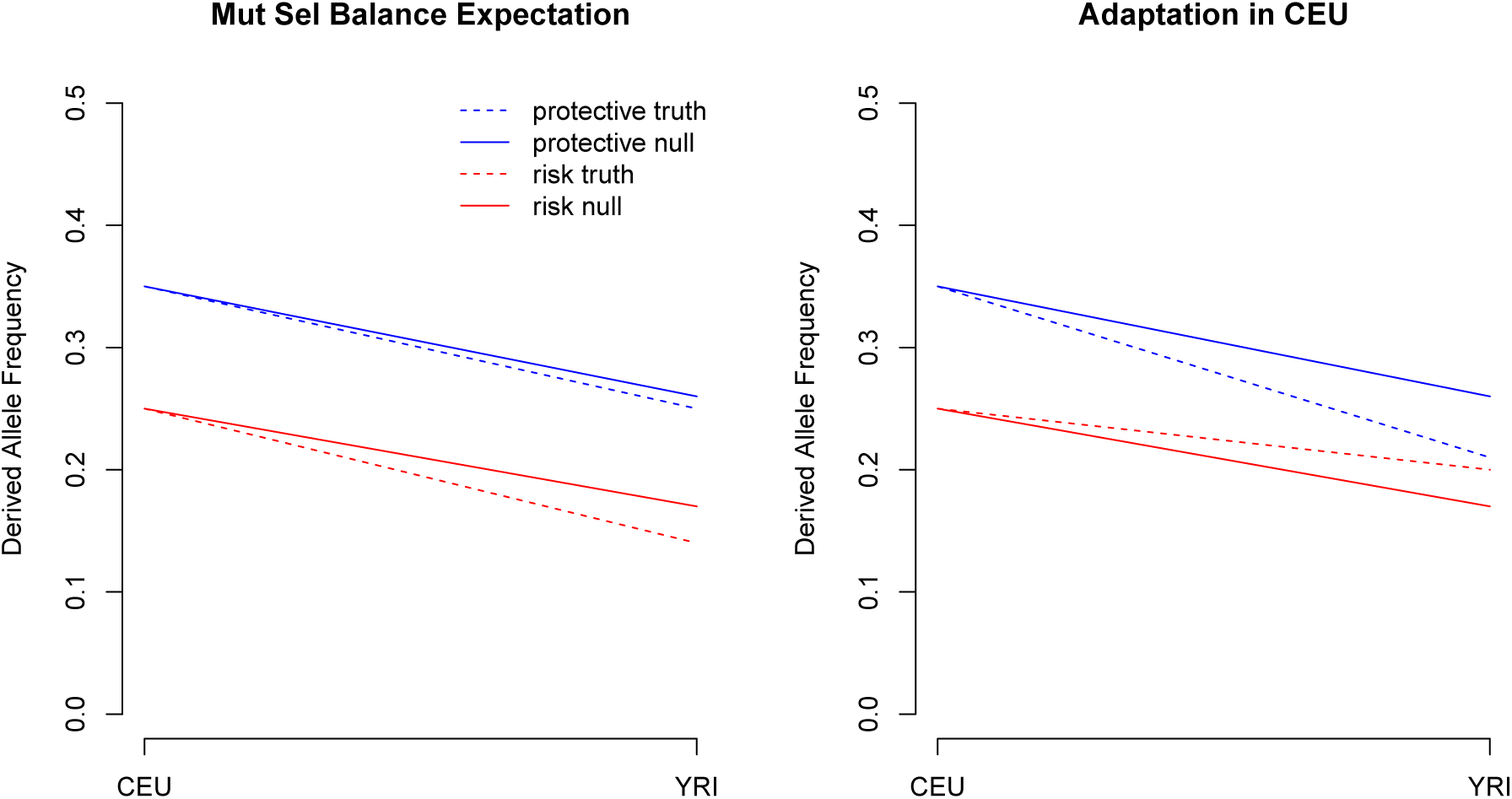
Two hypothetical patterns in the distribution of protective and risk alleles expected for disease traits which are *a priori* deleterious ancestral groups who had diverged in polygenic height scores.

To further explore this we looked at the decomposition of European populations by ancient ancestries. Haak *e*t al.^43^ has previously shown that most modern populations with European ancestry can be well described as a two-way mixture of WHG (modeled by Loschbour ancestry) and early Neolithic ancestry (modeled by LBK EN ancestry), followed a third wave of ancestry from the Yamnaya Steppe populations. For each of the European HO panel population samples we extracted the proportions of WHG, early Neolithic ancestry, and Yamnaya ancestry from Haak *e*t al.^43^ (these are reproduced in Figure S9). In Figure S10A-C we plot the proportion of each of these ancestries against the polygenic height for each modern European HO population (calculated at our subset of 724 SNPs). For comparison to these modern values we also plot, as horizontal lines, the mean polygenic height of the CEM (the analysis cluster containing LBK EN), the WHG (which contains the Loschbour individual), and the STP (which contains the Yamnaya samples).

Much of the variation in ancestry is along the Yamnaya-Early Neolithic axis, with the pearson correlation between Yamnaya and Early Neolithic ancestry being −0.92 across populations. There is somewhat less variation in WHG ancestry among modern European populations, WHG ancestry is corelated with Yamnaya ancestry (a pearson correlation of 0.60) and negatively correlated with Early Neolithic ancestry (a pearson correlation of −0.85). Modern European polygenic height scores are strongly correlated with the Yamnaya-Early Neolithic axis (e.g. pearson correlation of Yamnaya ancestry and polygenic height score is 0.69, p-value 0.0002, see also Mathieson^89^). To see how well Norwegian Lithuanian Estonian Icelandic Scottish Czech Belarusian Hungarian Ukrainian English Orcadian French_South Croatian French Spanish_North Bulgarian Tuscan Basque Bergamo Spanish Greek Albanian Sardinian modern European polygenic height scores could be predicted just on the basis of ancient polygenic height scores alone, we predicted each modern population (i) based on its admixture proportions from the Yamnaya (f_i,Y am_), Early Neolithic (f_i,EN_), and Western Hunter Gatherer (f_i,W HG_) as

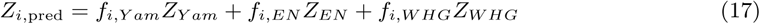

where for Z_Y am_, Z_EN_, and Z_W HG_ we used the polygenic score of the STP, CEM, and WHG respectively (calcuated over our 724 SNPs, calculated as described in main text). In Figure S10 we show these predictions plotted against the observed polygenic height scores (for the same SNP set). Overall the prediction works reasonable well. The prediction captures much of the overall height of the European polygenic scores, and the variation among populations. This suggests that the overall level of polygenic scores in modern Europeans is mostly attributable to high polygenic scores of WHG and STP populations, and that variation in polygenic heights is well explained by the differential contributions of early Neolithic, Western Hunter-Gatherers, and Early Neolithic populations to the ancestry of modern populations. While a reasonable fit, many of the observed values fall somewhat above their predictions. This is consistent with Europe-wide selection for increased polygenic height scores since the Yamnaya expansion, or (perhaps more realistically) that one of our source heights is mis-estimated by the ancient proxy. For example, perhaps the Yamnaya population who contributed ancestry to Europeans had a higher polygenic height score than that of our proxy sample.

**Figure S9:**
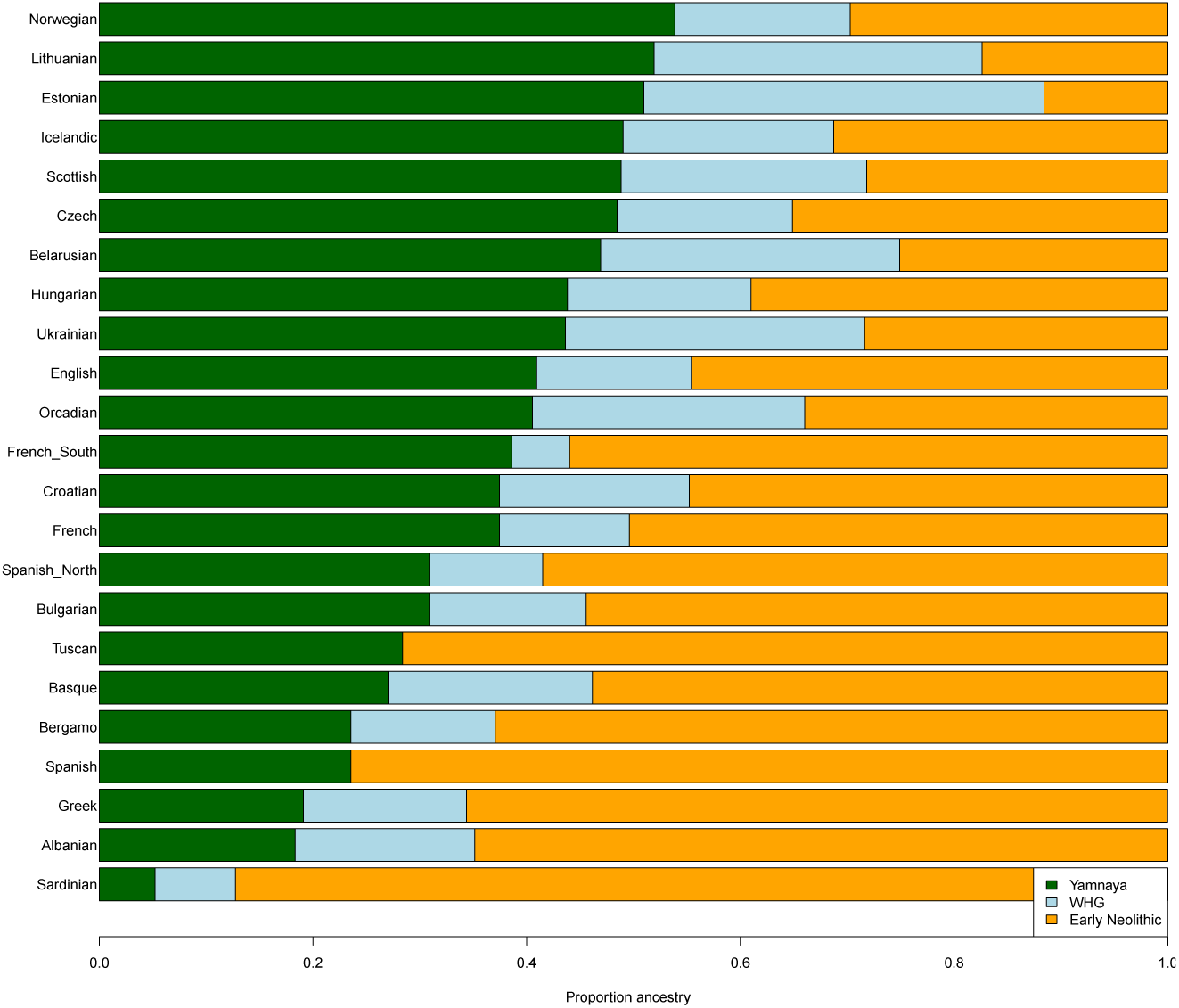
Modern European Admixture proportion estimates taken from Haak *e*t al.^43^

**Figure S10:**
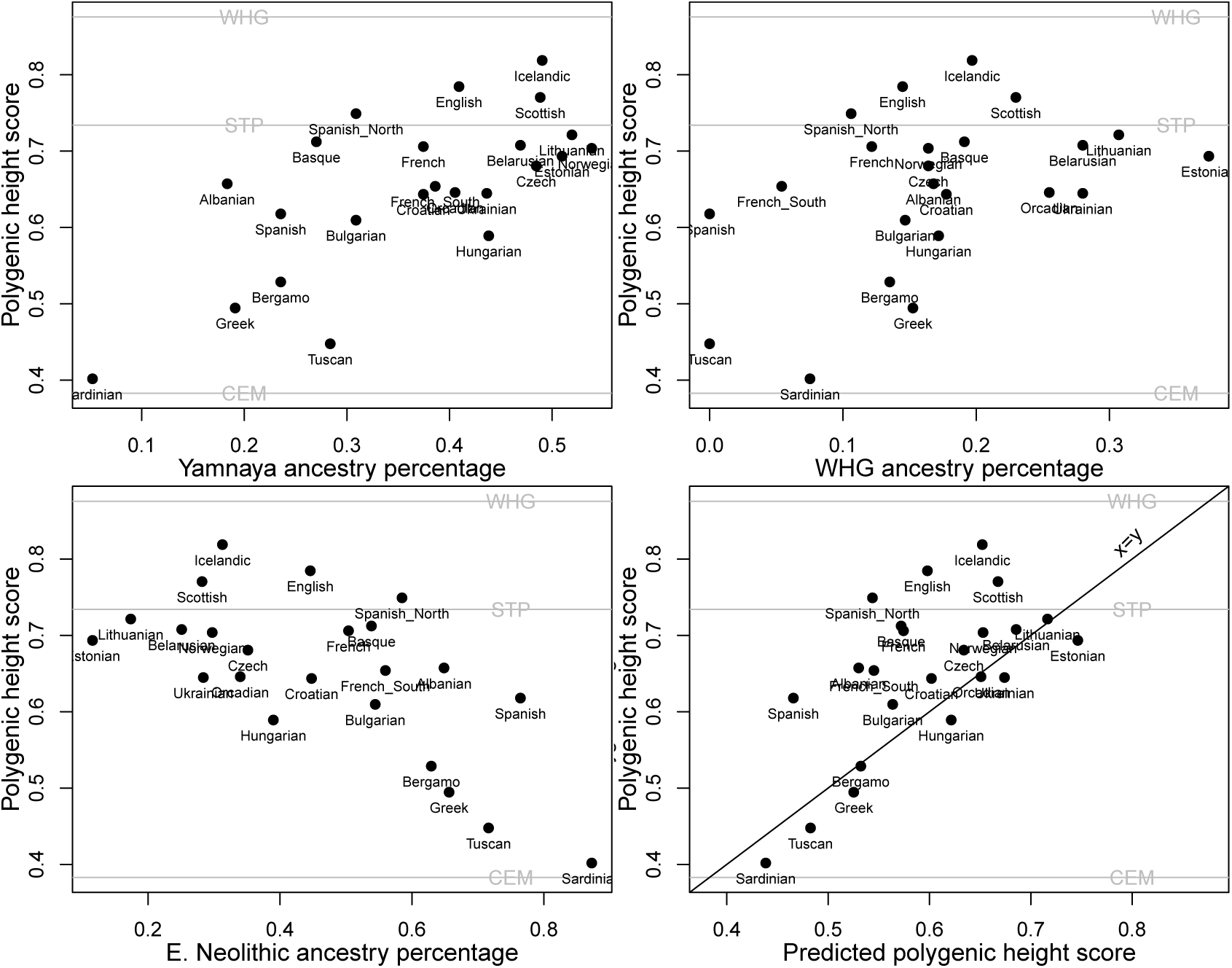
**A-C** Polygenic height scores for modern European populations plotted against their ancestry proportions for Yamnaya Steppe, WHG, and early Neolithic ancestry. The grey horizontal lines give the polygenic height scores for the WHG, STP, and CEM (central European Early and Middle Neolithic) population samples. **D) Polygenic height scores across modern European populations plotted againt a prediction for the scores based on ancient populations.**

### S1.6 Two Trait tests

#### S1.6.1 Conditional Tests

If 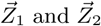 are vectors of polygenic scores for two different traits constructed according to equation (1), and the matrix 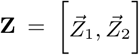 contains these vectors as columns, then under neutrality the distribution of **Z** is approximately matrix variate normal

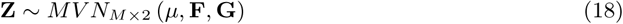

where the matrix µ contains the trait specific means, **F** gives the covariance structure among rows as in the single trait model, while **G** gives the covariance structure among columns. The matrix **G** is the canonical among trait genetic covariance matrix, or the "G matrix" of multivariate quantitative genetics,^47^ estimated for a population ancestral to all populations in the sample. The diagonal elements of this matrix are given by the V_A_ parameters from above in the single trait model, calculated independently for each trait. Off diagonal elements correspond to the additive genetic covariance between the two traits. In the case where some loci contribute to both traits, and all loci are approximately unlinked, these genetic covariances can be calculated as

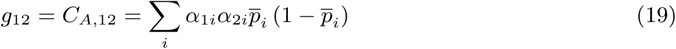

where α_1*i*_ and α_2*i*_ are the effects of locus i on traits 1 and 2 respectively. Given this null model for the joint distribution of the two traits, we can construct a conditional model for the distribution of trait 1, given values observed for trait 2, as

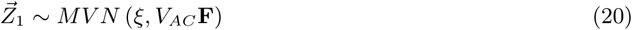

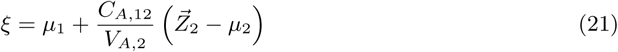

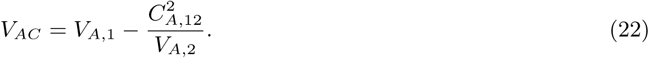

In the case where loci contributing to the two different traits are not unlinked, equation (19) is not an appropriate expression for the additive genetic covariance among traits, as it will depend also on the structure of linkage disequilibrium among sites. Because we ascertain SNPs independently for the two different traits, we expect that we will frequently have cases where two SNPs affecting two different traits within the same block are in linkage disequilibrium with one another, and therefore do not drift or respond to selection independently. To deal with this issue, we represent the genetic covariance among populations with a general form

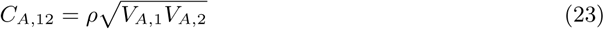

where ρ represents the genetic correlation between the two sets of polygenic scores. Further, we treat this genetic correlation parameter as an unknown, and also allow for one choice of genetic correlation parameter (ρ1) to describe the response of the mean, and a second correlation parameter to describe effects on the variance. The final two trait conditional model is then

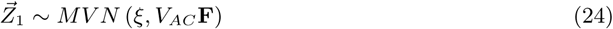

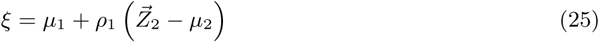

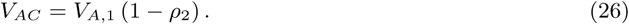

We calculate a conditional version of our test statistic:

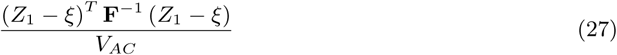

for a two dimensional grid of values for both ρ1 and ρ2 ranging from −1 to 1 and report the most conservative test. This procedure is overconservative, and therefore any trait which cannot be adequately explained explained as a response to some other trait under this framework is assume to have experienced an independent response to selection.

#### S1.6.2 Alternate Ascertainment Tests

As a second test, we constructed polygenic scores for each of HIP, WHR, IHC, and WC (which we denote with the prefix hsnp) using the subset of height SNPs for which an effect size estimate was available (about 1300 in each case), and then applied our test to these polygenic scores. Both hsnpHIP and hsnpWC show strong evidence of selection (p = 1.16 × 10^−14^ and p = 3.16 × 10^−6^ respectively), while hsnpIHC and hsnpWHR each shows no convincing signal (p = 0.07 and p = 0.28 respectively).

Combined with our results from the conditional model described above, this suggests that selection on height (or something tightly correlated to it) has plausibly impacted the genetic basis of HIP, WC, and probably IHC (though this alternate ascertainment test does not show evidence of, the conditional test does, and the moderate genetic correlation between them suggests it is likely). However, the conditional test cannot fully explain patterns observed for IHC and WHR given height, suggesting the action of independent selection. It is also conspicuous that we observe a stronger signal of selection for hsnpWC that for WC itself, and that the two are negatively correlated after accounting for population structure (r = −0.24, p = 9.5 × 10^−4^; though the presence of structure actually serves to mask the correlation; see Figure S11). Given the moderate *positive* genetic correlation between height and WC within populations, this negative correlation is surprising, and suggests that WC has been impacted by selection independent of height. WHR seems the most plausible candidate of the phenotypes included in our study.

**Figure S11:**
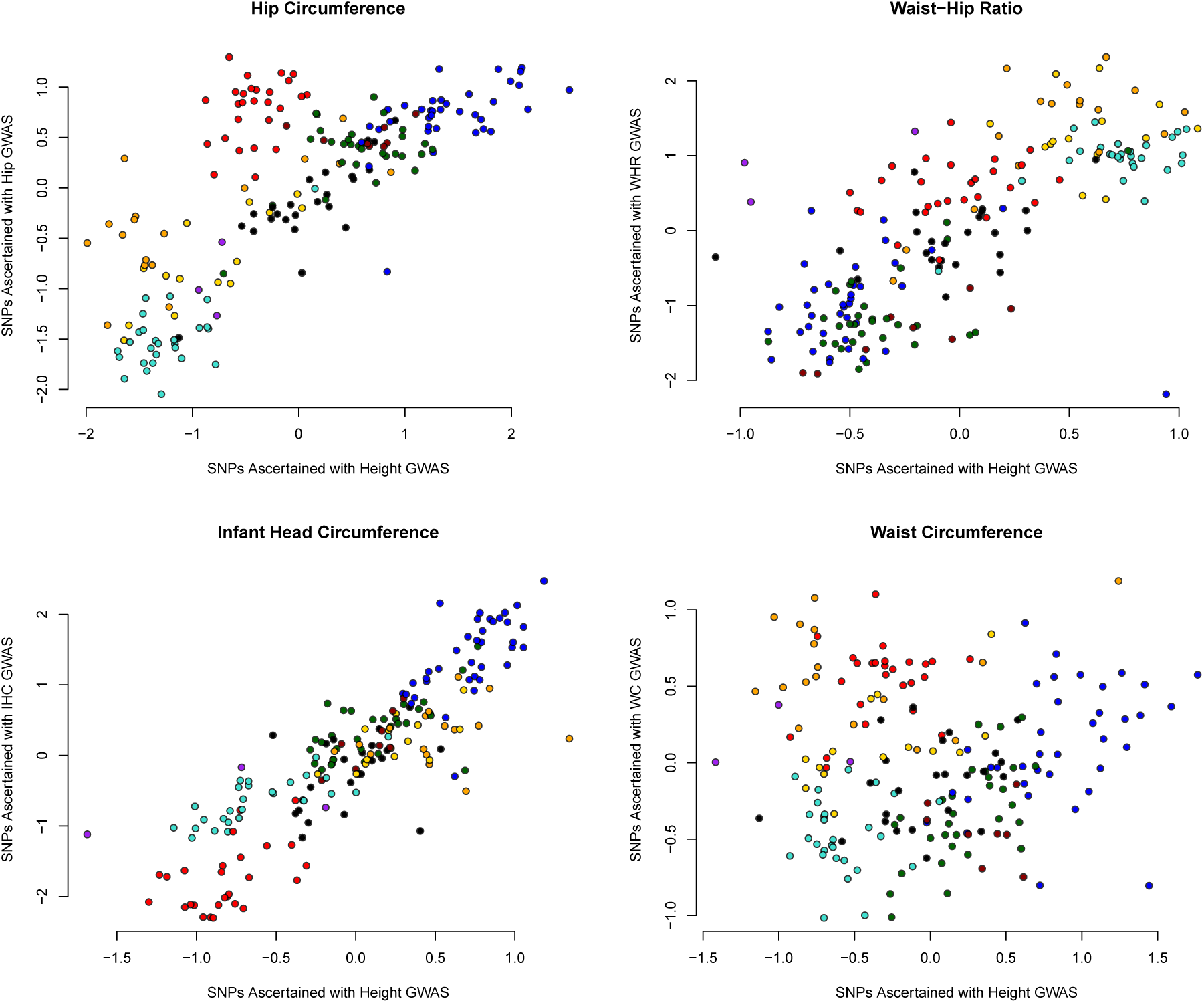
Polygenic scores for HIP, WHR, IHC, and WC plotted against scores computed from SNPs ascertained on the basis of association with height using the nominal effect size for the focal trait.

### S1.7 Allometry

Many phenotypes have allometric relationships,^90, 91^ e.g. waist circumference does not increase linearly with the height across individuals within a population measured at the same growth stage(s) (termed individual or static allometry^92^). One natural concern therefore is that in ruling out that selection signal for (say) waist circumference can be completely explained by a correlated response to selection on height, we have not dealt with the allometric relationship among the phenotypes across populations.^93^ Our linear prediction of (say) waist-circumference conditional the observed polygenic scores for height in theory will not capture the non-linear relationship between the two phenotypes, and our claim of an independent signals in height and waist circumference might be suspect. The easiest way to see that this may not be too much of a concern is to note that while our results are significant the differences in polygenic score we see among populations all correspond to relatively small shifts in phenotype. Therefore, even though the phenotypes have an allometric relationship this will be approximately linear over the scale we are looking over and so well accounted for by our multivariate approach. To demonstrate this more thoroughly we convert our polygenic scores to a phenotype-scale, multiplying by the standard deviation of the phenotype and adding the mean phenotype, and plot two phenotypes on a log-log axes (e.g. height to waist circumference top panel in Figure S12), noting again that we do not view these as accurate phenotypic predictions). Confirming the idea that our deviations are small in phenotypic space, the log-log (base e) axes gives a very similar picture to the linear axes (top and bottom panel of Figure S12) suggesting that allometric scaling is not a concern.

We can however offer a more general response to the concerns about allometry, based on the fact that we construct our polygenic scores based only on the contribution of each locus to the additive variance (and no higher variance components). An observed allometric relationship between the underlying genetic phenotypes implies non-additive genetic covariance among the phenotypes (see e.g. Rice (1998)^94^). To see this note that an allele with a fixed additive effect on height has a variable additive effect on waist circumference that depends on the distribution of allele frequencies at all of the height loci in the genome. For example, as WC has a positive allometric relationship with height, in a population with a high frequency of short height alleles the effect of our fixed height allele on waist circumference is smaller than when the population consists of many tall height alleles. Therefore, an allometric relationship between a pair of genetic phenotypes (among individuals or populations) implies that there is dominance and epistatic genetic covariance among the loci contributing to our traits (and some amount of epistatic variance in one or both traits). However, our polygenic scores are strictly based on the additive effect sizes, therefore they can not capture these higher order covariance components. As we are missing these higher order covariance terms our polygenic scores can fail to predict the phenotypes correctly due to allometry, but importantly also there can only be a linear relationship between our polygenic scores. Therefore our inferences of independent selection on the polygenic scores of multiple phenotypes cannot be confounded by allometry between phenotypes.

### S1.8 Height and the "thermoregulatory hypothesis"

To study the "thermoregulatory hypothesis", Ruff (1994)^2^ proposed a simple geometric model of body shape, where individuals are imagined as cylinders. The height of the cylinder is given by the individual's height (h), and the cylinder's circumference (C) by waist or hip circumference (note that Ruff modeled the diameter of the cylinder by bi-iliac breadth). Based on this we can calculate the surface area as *S* = *hC* +4π (*C*/2π)^2^; the volume as *V* = 2π (*C*/2π)^2^ h; and their ratio *V /S*. Based on these relationships we can compute the effect size of a SNP on surface, volume, and volume/surface ratio (α_Surf_, α_Vol_, α_Vol/Surf_) based on the effect of a SNP on height and waist circumference (α_Height_, α_Waist_), using a first order Taylor series approximation.^95^

We applied this procedure to all of our height SNPs, using the estimates of their effects on height and waist circumference, and in Figure S13 we plot their effect on cylinder surface, volume, and volume/surface. Using the effect of height SNPs effect on hip circumference yields qualitatively similar results. Increasing height does indeed have a positive effect on V/S ratio. However, as is to be expected from the weak direct dependence of a cylinder's V/S ratio on height, this is almost entirely driven by the correlated effect of SNPs on (waist or hip) circumference. As we cannot explain our height signal as a correlated effect of selection on hip or waist circumference, it seems unlikely that selection on height is mediated only through selection on V/S ratio.

### S1.9 Allen's Rule and Sitting Height Ratio

To explore the effect of selection on leg and torso length we took our set of height SNPs and extracted their effect on sitting height ratio (SHR) from a SHR GWAS^59^). We found no net relationship between an allele's effect on height and it's effect on SHR (Spearman's ρ = −0.010, p-value = 0.66, Figure S14A). This suggests that height SNPs act both on leg and head+torso length, as well some SNPs that affect each trait somewhat separately, see Chan et al.^59^ for more discussion. We compute a polygenic score from the effects of these height SNPs on SHR and found no signal of over-dispersion (Q_X_ p-value=0.19). This suggests that while selection has driven divergence in polygenic height scores, we do not have evidence that this selection has driven differences in the body-proportions of height.

To extend this observation we used the effect size of height SNPs on height and SHR ratio to estimate the unobserved effect of height SNPs on leg and torso+head height. We denoted the unknown effect sizes of the l^th^ SNP on leg and torso+head length by α_Ll_ and l_Tl_. The additive effect sizes of a SNPs on height and SHR are α_*Hl*_ and α_*Rl*_ respectively. While we will denote the standard errors of these effect sizes by σ_H,l_ and σ_Ll_. We model the additive height effect size as

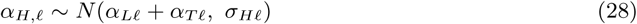

and, by a first-order Taylor series approximation, the SHR ratio effect size is modeled as

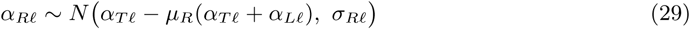

where µ_R_ is the population mean sitting height ratio (µ_R_ = 0.52).^59^ We assume that the parameter pair (α_Ll_, α_Tl_) ~ *N* (0, Ω), over all our loci, where Ω is the 2 × 2 variance-covariance matrix between leg and torso+head effect sizes. We place hyper priors on the diagonal elements of Ω (Ω_i,i_ ~ Cauchy(0, 1), Ω_i,i_ ≥ 0) and on the covariance 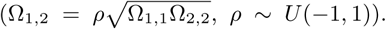 We then estimate α_Ll_ and α_Tl_ over all loci. In Figure S14B-F we plot α_Ll_ and α_Tl_ against α_Hl_ and α_Rl_ and each other.

**Figure S12:**
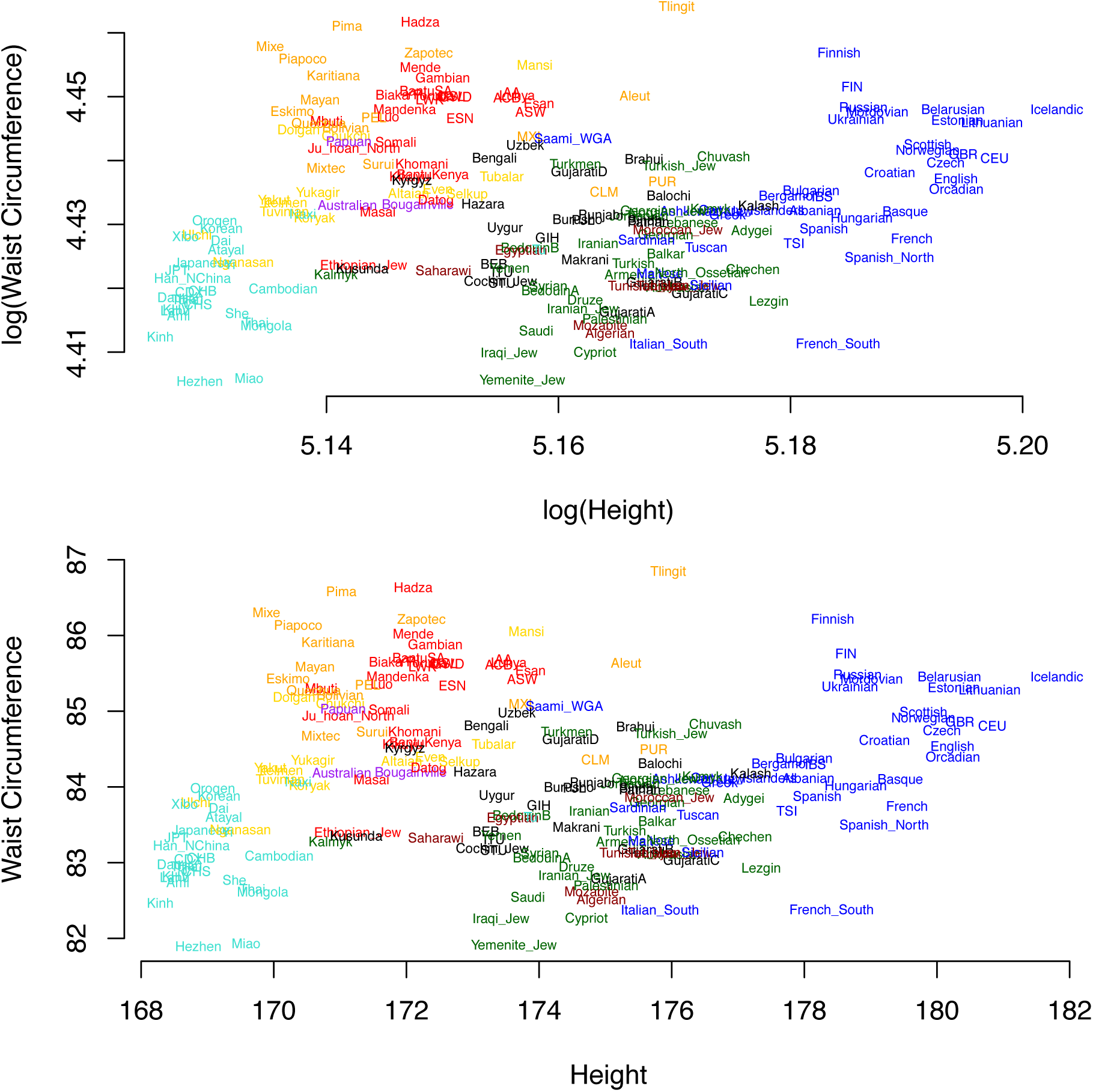
Relationship between polygenic scores, placed on phenotypic scale, for height and waist circumference on a log-log axis (top) and standard axes (bottom)

**Figure S13:**
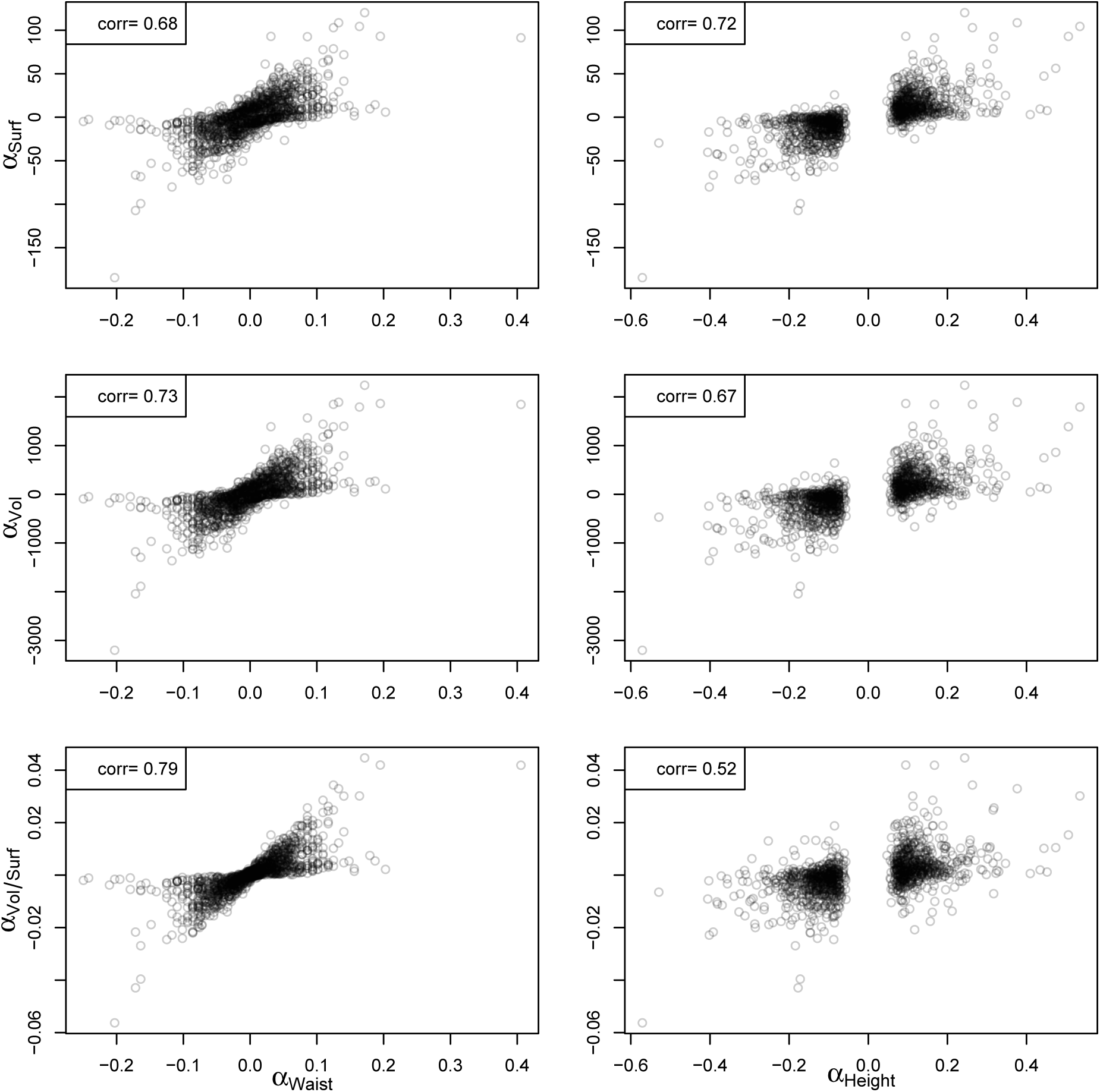
Relationship between the effect size of a SNP allele on height and waist circumference and the predicted effect of a SNP on surface area and volume. The pearson correlation coeffcient between the two variables is shown in the top right corner of each plot.

Using these estimates of effect sizes for leg and torso+head length we obtained Q_X_ p-values of 4.0 × 10^−32^ and 1.5 × 10^−31^ respectively in our total sample. Thus both leg length and torso+head length show a strong signal of responding to selection on height.

**Figure S14:**
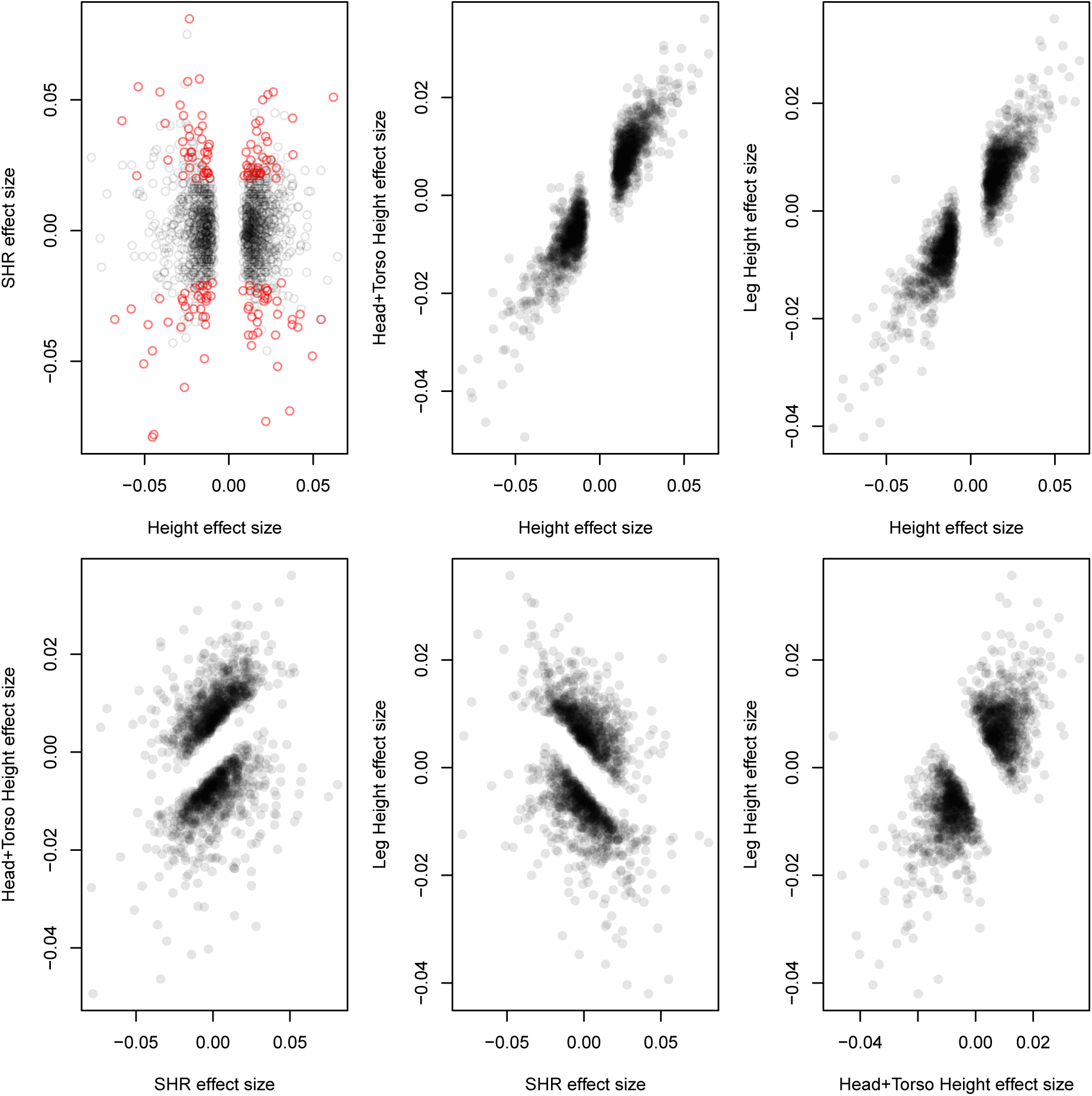
Plots of SNP effect sizes for height (α_*Hl*_) and SHR (α_*Rl*_) and the estimated effect sizes for leg (α_*Ll*_) and torso+head length (α_*Tl*_ α_*Tl*_) plotted against each other over SNPs. See Section S1.9 for more details. In the top left panel SNPs that have a significant effect on SHR (*p <* 0.05 are colored orange.)

